# Conserved cell state dynamics reveal targetable resistance patterns in ovarian high-grade serous carcinoma

**DOI:** 10.1101/2025.06.13.659489

**Authors:** Anna Pirttikoski, Laura Gall-Mas, Wojciech Senkowski, David Fontaneda-Arenas, Matias Marin Falco, Erdogan Pekcan Erkan, Johanna Hynninen, Krister Wennerberg, Anna Vähärautio

**Affiliations:** Research Program in Systems Oncology, Research Programs Unit, Faculty of Medicine, University of Helsinki, Helsinki, Finland; Biotech Research and Innovation Center, University of Copenhagen, Copenhagen, Denmark; Department of Obstetrics and Gynecology, University of Turku and Turku University Hospital, Turku, Finland; Foundation for the Finnish Cancer Institute, Helsinki, Finland

## Abstract

Core homeostatic programs of tissues, reflected in gene expression modules, can persist through oncogenesis. To reveal how cell states of normal fallopian tube epithelia (FTE) transform into intra-tumoral heterogeneity in ovarian high-grade serous carcinoma (HGSC), we applied a continuous, multi-state approach on single-cell transcriptomes of treatment-naive tumors (n=50) and normal FTE (n=14). We found the gene modules conserved from normal FTE to cancer to be more diverse than those altered during tumorigenesis, wherein the pseudotime-late TNFα/NF-κB–associated module was linked to poor prognosis. The modules showed biases in subclonal structures and homologous-recombination deficiency, suggesting that genetic drivers employ existing epithelial programs for oncogenic phenotypes. Finally, organoid experiments validated actionable non-genetic drivers of these cell states, allowing chemosensitization by modulating apoptosis and NF-κB pathways. In summary, we identified prognostic, continuous cell states that reconstitute fallopian tube dynamics in HGSC and offer actionable targets to improve platinum response.

## Introduction

Transcriptional heterogeneity in cancer is increasingly acknowledged as a key factor in tumor progression, metastasis, and therapy resistance (1–3). Ovarian high-grade serous carcinoma (HGSC) is characterized by late detection, frequent platinum resistance, and a high mortality rate. Most patients relapse, leading to refractory disease with a 5-year survival rate of 43% (4). Overall survival (OS) has only improved among patients with homologous recombination-deficient tumors benefiting from poly (ADP-ribose) polymerase (PARP) inhibitors (5). From a genomic perspective, HGSC is a cancer predominantly driven by copy number alterations and has nearly universal occurrence of *TP53* mutations (6,7). HGSC shows high inter- and intratumoral heterogeneity driven by genetic, epigenetic, and transcriptional alterations (8,9). We have already found that diverse, pre-existing or therapy-induced transcriptomic states of HGSC cells are associated with treatment responses, playing a key role in drug resistance (10, 11).

Recent studies have given strong evidence that HGSC primarily originates from the fallopian tube epithelium (FTE) based on transcriptomic (12) and evolutionary studies (13) as well as mouse models (14). Lesions with identical *TP53* mutations to fallopian tube have been identified with the HGSC patients, and this p53 signature develops eventually to HGSC through serous tubal intraepithelial carcinoma (STIC) (15). Molecular analysis further supports this, showing that HGSC transcriptomes share more similarities with FTE than ovarian surface epithelium (OSE) (16). Previously, a comparative single-cell RNA sequencing (scRNA-seq) analysis has found a connection between diverse secretory cell populations from FTE and HGSC subtypes, which also highlights how phenotypic heterogeneity in cancer cells is inherited from their cell of origin (12). In addition, FTE scRNA-seq analyses have provided functional insights into HGSC through deconvolution analysis of precursor states (17), molecular subtype stratification based on secretory cell markers (18), and the exploration of GWAS-derived risk genes (19). Nonetheless, these studies face limitations, including a restricted number of postmenopausal donors and the unavailability of single-cell transcriptomic data from HGSC tissues, which precludes direct high-resolution comparisons.

Cancer survival and tumor progression are dependent on phenotypic heterogeneity within cancer cells. Recent single-cell transcriptomic studies across various cancer types have revealed the coexistence and plasticity of diverse cell states among cancer cell populations (20, 21, 22). Cell states contain information about multiple aspects of cell identity, such as metabolic state, cell cycle phase and cell-specific molecular signatures. Therefore, an individual cell can express several functional gene modules in parallel to achieve a variety of phenotypic characteristics (23). Consequently, the state of cancer cells is likely shaped by the diverse expression of gene modules (24). There is evidence that most gene modules found from tumors are not exclusively expressed in cancer but are derivatives of those induced in normal physiological processes to maintain tissue homeostasis (25). These co-opted gene modules from the tissue of origin have been found to represent several biological pathways, such as stress (26) and interferon response (27), as well as differentiation processes (28). Overall, the remarkable plasticity of cancer cell states and their reliance on non-genetic mechanisms are underscored by the ability of individual cancer cells to reconstruct tumors that recapitulate the transcriptional programs of their cells of origin (20, 29).

To reveal how the multifaceted cell states of normal FTE are inherited -and distorted-into intra-tumoral heterogeneity in HGSC, we apply here a continuous, multi-state approach in single cell transcriptomes of treatment-naive tumors (n=50) and normal fallopian epithelia (n=14). This continuous gene module approach contrasts with previous studies that categorize intratumoral, chemotherapy driven (10) and normal epithelial heterogeneity (12, 17, 19) into discrete cell states, thus offering a more flexible and potentially more biologically realistic perspective. Furthermore, previous studies addressing FTE subpopulations have -due to limited numbers of HGSC samples analysed by scRNA-seq-projected the normal states onto bulk RNA-sequencing data (12, 17, 18), therefore lacking high-resolution alignment between heterogeneity in normal epithelia and cancer.

Our analysis revealed a pair of contrasting gene modules conserved from normal FTE to cancer having a prognostic significance in HGSC at a patient level. We discovered that a TNFα/NF-κB pathway-enriched state predicts poor clinical outcome, identified candidate compounds targeting this state, and demonstrated that its modulation influences chemosensitivity in matching patient derived-organoid (PDO) cultures.

## Results

### Inter-patient heterogeneity formed during HGSC tumorigenesis

We collected 52 tumor tissue samples from 50 treatment-naive patients with HGSC from several anatomical locations: peritoneum, omentum, mesentery, ovary, ascites, adnexa, or fallopian tube. We obtained 7 control samples from fallopian tube or fimbriae from healthy individuals who were postmenopausal women without earlier HGSC diagnosis. One of the control samples was from a patient having Li-Fraumeni syndrome with a germ-line mutation in the *TP53* gene. Detailed clinical information of the patient cohort is given in Table S1. In addition, we incorporated external scRNA-seq data from secretory epithelial cells derived from 7 postmenopausal donors (19) into our gene module construction to increase sample size and to improve robustness of our analysis.

We measured transcriptomes of cells using scRNA-seq (Fig. 1A) and identified malignant epithelial, secretory FTE, immune, and stromal cells (Fig. S1A-D, Table S2 and see Methods). Unlike stromal and immune cells which clustered together across different patients, cancer cells displayed a distinct expression pattern unique to each patient (Fig. S1D), consistent with earlier studies (8, 30). Secretory epithelial cells from FT grouped apart from cancer cells and showed a lower level of patient-specificity (Fig. 1B). This was evidenced by Louvain clustering, where on average, 96% of secretory cells from each FTE sample were clustered within the shared cluster. On the contrary, cancer clusters showed high patient-specificity when, on average, 84% of the cells within the cancer clusters originated from the same patient. Collectively, our data suggest that the relative homogeneity of gene expression in normal cells compared to cancer cells supports the hypothesis that inter-patient heterogeneity develops as a consequence of HGSC tumorigenesis.

**Fig. 1.**
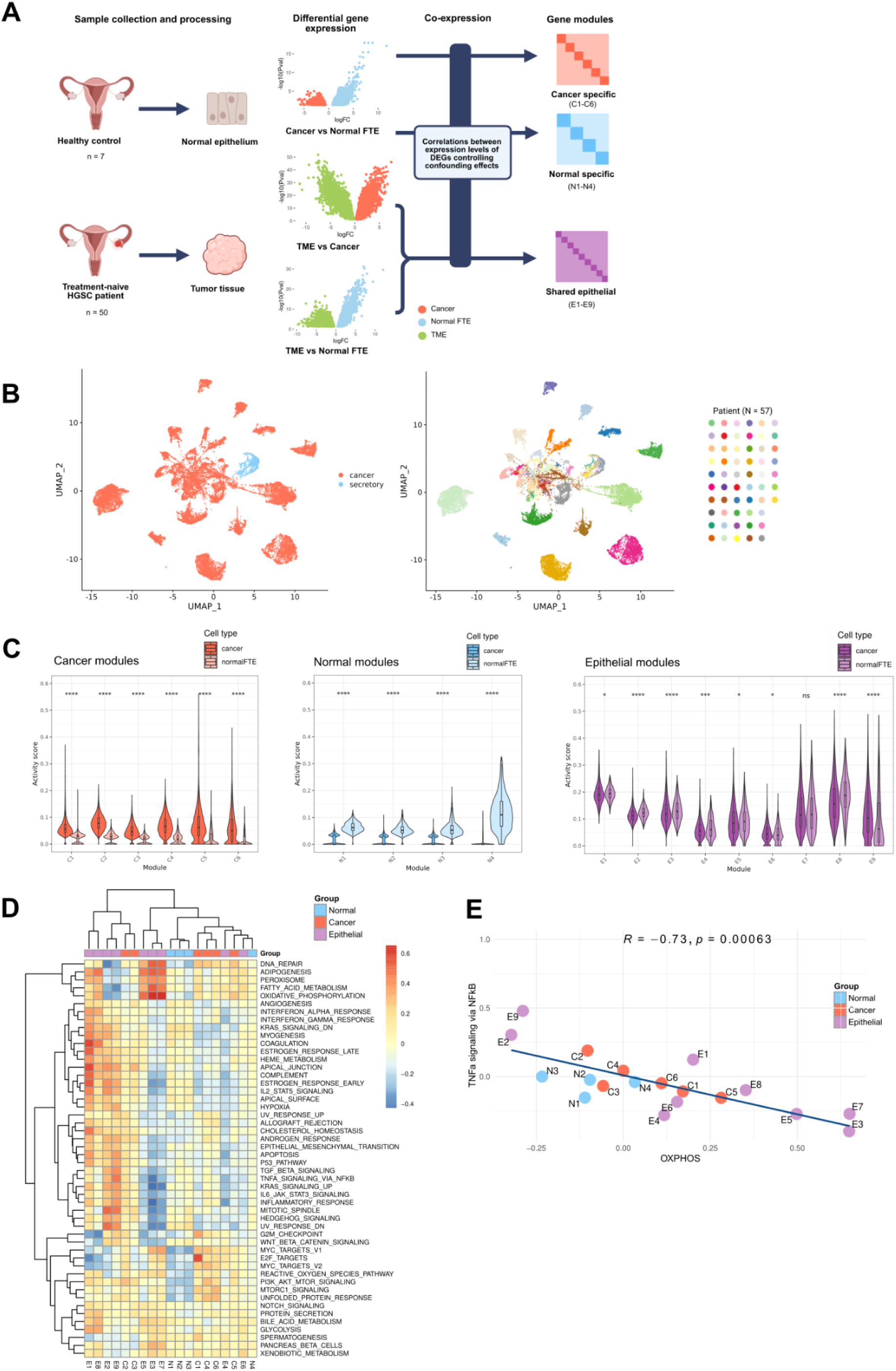
Overview of sample processing and the identification of gene modules and their characteristics in HGSOC. (**A**) Overview of the scRNA-seq data and gene module detection approach. Cancer and normal specific as well as shared epithelial genes were identified using pseudobulk DGE analysis where external FT secretory scRNA-seq data set were utilized. Co-expression between the DE genes were calculated with a tool controlling confounding effects. Hierarchical clustering based on WGCNA was applied to identify gene modules based on co-expression patterns. (**B**) UMAP of 31 487 epithelial cells colored by the sample origin (tumor or FTE) (left) and by the patient (right). (**C**) Violin plots showing module activities in cancer vs secretory FTE cells using single-cell gene signature scoring method UCell. (**D**) Correlation heatmap between gene modules and hallmark gene set collection from MSigDB calculated at a single-cell level of cancer cells (n = 8 786) using UCell for the calculation of signature activity. (**E**) Correlation plot of TNFα signaling via NF-κB and OXPHOS across all the gene modules (n = 19) in cancer cells, indicating an inverse relationship. Data points are colored by module group.

### Oxidative phosphorylation and TNFα-signaling define opposing, conserved gene modules

Transcriptional heterogeneity among cancer cells within a tumor has been found to be organized into non-discrete cell states where several gene programs can be active at the same time (24). Considering this complexity of cell states, we allowed an individual cell to express combinations of gene modules and used our unique gene module detection approach to characterize the heterogeneity of treatment-naive HGSC cells and normal secretory epithelia (23) (Fig. 1A). To capture population-level gene expression differences and ensure detection of lowly expressed genes, we conducted pseudobulk differential gene expression (DGE) analysis to identify distinct expression patterns between cancer and secretory FTE cells. As gene modules are generally not exclusively expressed in cancer cells but are often common to those activated in respective normal tissue (25), we also determined shared epithelial genes overexpressed in both cancer and FTE cells compared to cells of the tumor microenvironment (TME).

To identify cancer-specific, normal FTE-specific and conserved epithelial co-expressed gene modules that are present across the patients, we grouped the DEGs within each cell group based on their similar expression patterns using CS-CORE (31) and WGCNA (32) methods (see Methods). With this approach, we identified six cancer-specific (C1-C6), four normal-specific (N1-N4) and nine shared epithelial (E1-E9) co-expressed gene modules (Table S3). We compared the activity of these gene modules between cancerous and normal FTE cells using the UCell method (33). As expected, cancer-specific gene modules were significantly more active in cancer cells, while normal-specific modules were more active in FTE cells (*P* < 2e-16). Epithelial gene modules were more uniform between cancer and normal FT epithelial cells (Fig. 1C). We also compared module activities between the normal FTE sample from the patient with Li-Fraumeni syndrome and other normal FTE samples, observing significant differences particularly in normal module activities. In contrast, the Li-Fraumeni syndrome FTE sample exhibited more similar activity levels to the other control samples for cancer and epithelial modules (Fig. S2A). In terms of *TP53* expression, epithelial cells from the Li-Fraumeni syndrome sample exhibited lower expression levels compared to other normal FTE samples (*P* = 0.011, Fig. S2B). Taken together, despite reduced *TP53* expression, germ-line *TP53* mutated FTE exhibited transcriptional states more closely resembled those of normal FTE than cancer.

Independence of the gene modules were examined in both cancer and FT epithelial cells by calculating correlations between them at single-cell level. Mostly there was no association between the gene modules, with the median of Spearman’s correlation coefficients 0 in cancer (IQR: −0.12 to 0.13) and normal FTE cells (IQR: −0.04 to 0.17). In both cancer and FTE cells, the strongest negative correlation was between modules E3 and E9, and the strongest positive correlation between E3 and E7 (Fig. S2C).

We evaluated the biological roles of the gene modules by assessing how they correlate with the hallmark gene sets from the Molecular Signatures Database (MSigDB) (34). Modules E3, E5, E7 showed moderate to strong association with oxidative phosphorylation (OXPHOS) in both cancer (r between 0.50-0.65) and normal FTE cells (r between 0.54-0.57) (Fig. 1D, Fig. S2D), and were also connected to fatty acid metabolism, adipogenesis and DNA repair. Module E9 was the highest linked to inflammatory processes, such as TNFα signaling via NF-κB (r = 0.48 in cancer cells) and to inflammatory signaling in normal cells (r = 70), also showing strong association with TGF-β signaling (r = 0.42 in cancer) and Epithelial to mesenchymal transition (r = 0.70 in normal FTE cells). It is noteworthy that in normal FTE cells, cancer modules showed no association with any of the cancer hallmark gene sets, while normal modules exhibited largely negative associations (Fig. S2D). This suggests that the non-conserved gene modules reflect distinct features between cancer and normal FT epithelial cells. Moreover, we found an inverse relationship between TNFα signaling via NF-κB and OXPHOS based on their associations with the gene modules in cancer cells (r = −0.73, *P* = 0.00063) (Fig. 1E). In normal FTE cells, this relationship was not found (r = −0.28, *P* = 0.24) as normal-specific states showed low levels of both (Fig. S2E). In summary, we deconstructed the transcriptional heterogeneity of cancer cells using a gene module-based approach guided by the cell of origin, and identified an inverse relationship between OXPHOS and TNFα signaling under cancerous conditions.

### A pseudotime-late cell state is associated with poor prognosis in HGSC patients

To investigate the potential prognostic significance of the gene modules in treatment-naive tumors at the patient level, we analyzed deconvoluted bulk RNA-seq data and clinical information from 258 patients from The Cancer Genome Atlas (TCGA) cohort (7, 35). Kaplan-Meier survival analysis indicated that patients with module E7-high tumors at diagnosis had significantly longer OS time compared to patients with E7-low tumors (log-rank test, *P* = 0.0025; Fig. 2A). Conversely, patients with module E9-high tumors at diagnosis had significantly shorter OS time compared to patients with E9-low tumors (log-rank test, *P* = 0.0015; Fig. 2A).

**Fig. 2.**
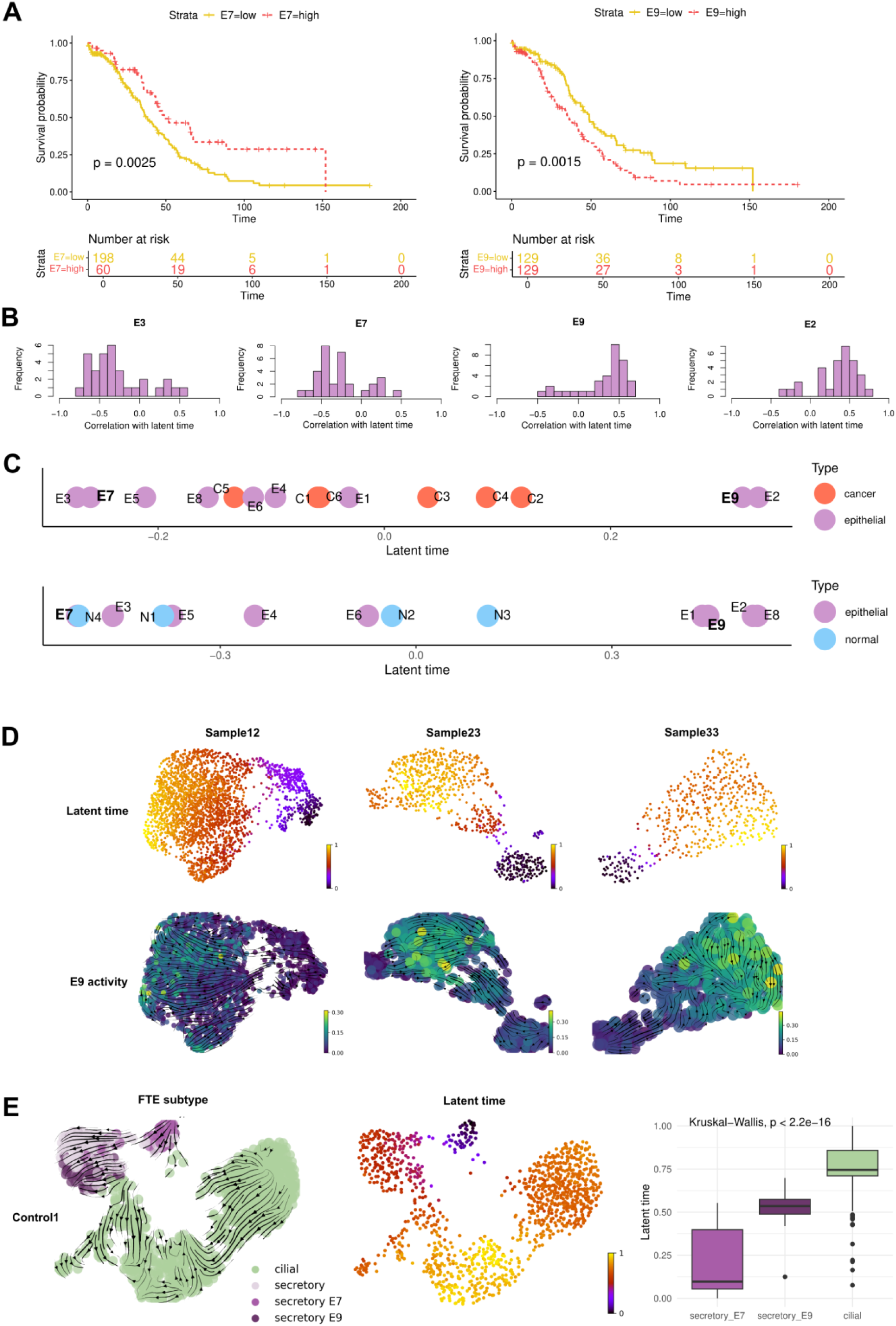
A developmentally late module is predictive of poor outcome in HGSC patients. (**A**) Kaplan-Meier curves on overall survival (OS) for module-high and module-low patients (log-rank test, *P* = 0.0025 for E7 module and *P* = 0.0015 for E9 module) in The Cancer Genome Atlas (TCGA) cohort. The number of patients at risk at each time point is listed below the survival curves. (**B**) Histograms showing distributions of cancer sample-specific correlations with a latent time for epithelial gene modules with the strongest average correlations. Only correlation coefficients having *P* < 0.05 are shown in histograms. (**C**) The average correlation between the gene modules and latent time scores across the cancer samples (above) and the FTE samples (below). Data points are colored by module group. (**D**) UMAPs of latent time scores calculated with CellRank and E9 module activity in three cancer samples. Latent time is represented on a scale from 0 to 1, with 0 indicating early and 1 indicating late state. Velocity vectors showed with E9 module activity. (**E**) UMAPs of control1 normal FTE sample from a patient having a germ-line *TP53* mutation. Cells are color-coded by FTE subtype, and the top 25% of cells with the highest activity in either E7 or E9 are also indicated (left), alongside their latent time scores (right). Velocity vectors and latent time are calculated with the CellRank algorithm. Box plots showing the differences in latent time in secretory cells with the highest activity in E7 or E9, and ciliated cells.

To characterize the identified gene modules in the context of dynamics, we analyzed their relationship to gene-shared latent time, as inferred through the RNA velocity approach (36, 37). Most gene modules showed no clear correlation with latent time, with average correlation coefficients between −0.2 and 0.2 (Fig. S3). However, modules E2 and E9 had a weak positive correlation (mean r = 0.33 and 0.32, respectively), while modules E3 and E7 had a weak negative correlation (mean r = −0.27 and −0.26, respectively) with latent time across the cancer samples (Fig. 2B-C). Similar patterns were seen in normal FTE samples, despite a small sample size (mean r = 0.52, 0.45, −0.46, and −0.52, respectively; Fig. 2C). Latent time and E9 activity were visualised in three cancer samples with UMAP embeddings to exemplify how the late cell population, as estimated by latent time, showed higher E9 activity (Fig. 2D).

To validate the ability of the RNA velocity framework to capture biologically meaningful dynamics, namely the FTE differentiation process, we applied it to secretory and ciliated epithelial cells in control samples. According to current scientific understanding, secretory cells give rise to ciliated cells in FTE (38). Consistent with this hypothesis, our analysis showed that secretory cells were positioned at earlier time points along the latent time axis, wherein secretory cells had lower latent time scores than ciliated cells in five of six samples analysed. Furthermore, secretory cells with high E7 module activity preceded those with high E9 activity along the latent time trajectory (Fig. 2E, Fig. S4). Combined, our results suggest that the dynamics of epithelial gene modules is conserved from normal FTE to cancer. Intriguingly, in cancer specimens, only the modules on extreme ends of pseudotime associate with survival, with the late E9 module linked to short OS, opposing the role of early, stem-like states in HGSC clinical resistance.

### Gene module activities reflect subclonal architecture and HRD status of the tumor

The activity of clinically relevant modules varied slightly between cancer and normal FTE as E9 showed higher activity in cancer cells compared to normal FTE cells (*P* < 2.2e-16) while no significant difference was observed for E7 (*P* = 0.81) (Fig. S5A). To evaluate variability in E7 and E9 module activities, we analyzed module scores at single-cell level and found differences in activation both between and within patients, with a mild inverse relationship across the patients (r = −0.25, *P* < 2.2e-16; Fig. S5B). To further investigate the differences between modules E7 and E9 at the single-cell level in cancer cells, we compared their activities to 14 signaling pathways using PROGENy inference from consensus signatures of perturbation experiments (39). Our analysis revealed that module E9 exhibited had similar activity patterns to MAPK, NF-κB and TNFα pathways (r = 0.43, 0.36 and 0.34, respectively), aligning with the Hallmark pathway analysis (Fig 1D), while module E7 correlated with Trail and Hypoxia pathways (r = 0.35 and 0.28, respectively) in the cancer cells (Fig. 3A, Fig. S5C). To assess the robustness of the identified modules, we evaluated module associations with signaling pathways using an external scRNA-seq dataset including HGSC cells from 41 treatment-naive patients (8). Consistent with our findings, E9 module was associated most with TGF-β, MAPK and NF-κB pathways (r = 0.21, 0,19 and 0.18, respectively), while module E7 showed negative associations, with NF-κB and TNFα pathways (r = −0.18 and −0.17, respectively; Fig. S5D). An inverse correlation between E7 and E9 was observed at single-cell level also in this dataset (r = −0.15, *P* < 2.2e-16).

**Fig. 3.**
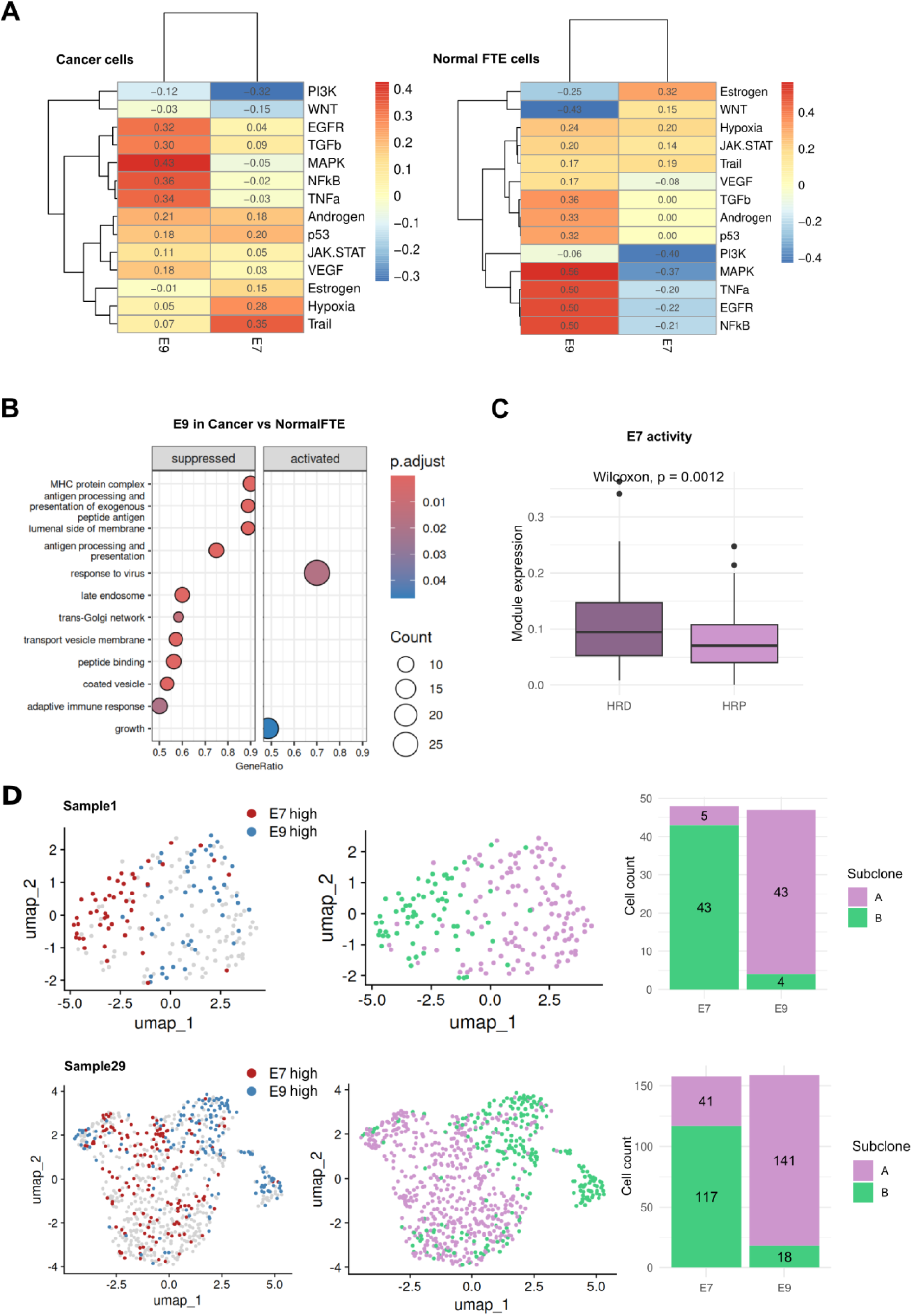
Pathways and genetic associations of the modules with clinical significance. (**A**) Correlation heatmaps of pathway activities using PROGENy with modules E7 and E9 in cancer cells (left) and normal FTE cells (right). (**B**) GSEA scores for Gene Ontology (GO) terms (with *P*.adjust < 0.05) were calculated between cancer cells and secretory FTE cells having the highest E9 module activity (top 25%) per sample. Genes originally differentially expressed between cancer and secretory FTE were filtered from the comparison (n = 234). (**C**) Box plot showing E7 module activity in homologous recombination deficient (HRD) (n = 138) and homologous recombination proficient (HRP) patients (n = 111) (Wilcoxon rank-sum test, *P* = 0.0012). (**D**) UMAPs of cells annotated with the highest module activity (left) and subclone identity (right) in two cancer samples. Stacked barplots showing the subclone distribution in E7- and E9-high cells.

In normal FTE cells, we also studied the pathway activity patterns of the modules, observing that the MAPK, NF-κB, TNFα, and EGFR pathways positively correlate with the E9 module, while negatively correlating with the E7 module (Fig. 3A). By examining expression differences in module E9 between cancerous and normal tissues, we found through gene set enrichment analysis (GSEA) that pathways related to antigen processing and presentation were significantly suppressed in E9-high cancer cells compared to E9-high normal cells (BH-adjusted *P* = 5.38e-05, NES = −2.75; Fig. 3B; see Methods)(40, 41). To delineate the source of the observed differences in antigen presentation, we compared its activity in the Li-Fraumeni syndrome sample to normal FTE and cancer, and observed that its activity profile was more similar to normal FTE (Fig. S5E). In addition, using the transcriptional signature contrasting E9 against E7 with GSEA, we found that transcription regulation, miRNA processing, and cell differentiation are activated in E9-high states, while cellular metabolism is suppressed (Fig. S5F). Epidermal cell differentiation was also upregulated in the E9-high state (BH-adjusted *P* = 2.48e-03, NES = 2.64), along with Notch and Wnt pathways, both linked to epidermal differentiation and cancer (Fig. S5G) (42).

Given that homologous recombination deficiency (HRD) is a key tumor characteristic predicting clinical response to PARP inhibitors and carboplatin therapy in HGSC, we analyzed the differences in E7 and E9 module activities in HRD and homologous recombination proficient (HRP) patients in TCGA cohort (43). We found that E7 module activity was significantly higher in HRD patients compared to HRP patients (*P* = 0.0012) whereas there was no difference in the E9 module activity (*P* = 0.78; Fig. 3C and Fig. S6A). We also included HRD status and patient’s age in multivariate Cox regression analysis and found that E7 and E9 modules were significantly associated with OS (*P* = 0.024 and 0.014, respectively) independently of the effects of HRD status or age (Fig. S6B). As *BRCA1* and *BRCA2* genes play a crucial role in HR DNA repair pathway, we validated the HRD association of module E7 using BRCAness transcriptional signature from an isogenic pair of parental BRCA1 mutated and wild-type (wt) BRCA1 restored COV362 ovarian cancer cells we created earlier (11). These results suggested that BRCA1 mutation related genes are upregulated (BH-adjusted *P* = 0.0016, NES = 1.42) and BRCA wt enriched genes downregulated in the E7-high state (BH-adjusted *P* = 2e-10, NES = −3.38).

Given HGSC’s genomic instability and copy number-driven nature, we assessed if frequently observed amplifications impact module activity. Using driver-amplified genes from Smith et al., we compared module activities in TCGA patients with amplified versus neutral copy number status (see Methods)(44). We found that E9 was more highly expressed in patients with *CCND3* amplification (*P* = 0.02), while E7 was lower expressed in patients with *AKT2* amplification (*P* = 0.045; Fig. S7A). However, these differences lost significance after *P*-value correction, likely due to the small sample size, and hence remain inconclusive (Table S4).

To evaluate subclonal enrichment of gene modules regardless of specific genomic aberrations, we used subclonal copy number variation (CNV) profiles of each patient inferred from scRNA-seq data (see Methods)(45). Across all patients and subclones, there was a significant link between subclone groups and module activity (*P* = 1.75e-264). In individual patients, 22 of 30 showed a significant association between module annotation and subclone identity after FDR-adjustment (Fig. 3D and Fig. S7B). For comparison, random cell assignments showed no association with subclone identity (Fig. S7B). Altogether, these results highlight that heritable differences influence the levels of these modules, and therefore the continuous states exhibit significant subclonal and HRD-associated biases despite lacking a clear genetic driver.

### Modulating chemoresponse by unbiased targeting of prognosis-linked gene modules in PDOs

To identify potential driver genes that contribute to the transition of cells into the poor prognosis E9 state, we employed data from Perturb-seq, a technique that associates pooled genetic perturbations introduced by CRISPRi with their transcriptional responses at the single-cell level (46). To allow more robust targeting, we expanded the gene signatures from restricted, epithelial-specific gene modules to genome-wide ordered gene lists by comparing cells with high module activity to those with low or inactive module states. Since module activities in the cells follow a skewed distribution without a clear threshold, we defined module-high cells as the top 25% of cells with the highest module activity scores in each patient (Fig. S8A). We also tested different thresholds and found that the core signal was robust against changing the threshold (Fig. S8B).

To identify candidate genes driving prognostic cell states, we compared E9 and E7 module signatures with transcriptional profiles resulting from gene perturbations, focusing on knockdowns negatively correlated with E9 and positively with E7 (Fig. 4A). *CCDC6* emerged as a top hit (E7: r = 0.23, adjusted P = 4.35e-85; E9: r = −0.19, adjusted P = 9.09e-58), showing significant expression differences between cancer and TME cells (P = 1.1e-09; Fig. 4B). *CCDC6* is involved in regulating DNA damage response, cell cycle control, and apoptosis, processes active in E9-high state (NES between 1.45-2.51; Fig. S8C). *ATP2A2*, negatively correlated with E9 (r = −0.10, adjusted P = 3.95e-18), is higher expressed in cancer cells than normal FTE or TME cells (P = 0.0061 and P = 1.4e-05; Fig. 4B), regulating calcium transport, with related processes active in E9-high states (NES 1.63-2.12; Fig. S8C). Beside these genes, we identified genes *LSM12* and *USP9X* as being associated with E7 (r = 0.15, adjusted *P* = 3.56e-36 and r = 0.11, adjusted *P* = 1.05e-23; respectively) and E9 (r = −0.17, adjusted *P* = 4.87e-48 and r = 0.11, adjusted *P* =1.87e-25; respectively) module signatures; however, their expression did not differ markedly between cancer cells and normal FTE cells or cells from the TME (Fig. 4B). *LSM12* has been found to play a role in activating Wnt signaling and promoting cancer progression (47), as well as participating in the cellular response to oxidative stress-induced DNA damage (48), which we also observed to be upregulated in the E9-high state (BH-adjusted *P* = 0.01, NES = 1.63). *USP9X* is a deubiquitinating enzyme and has been shown to promote metastasis and chemoresistance in various cancers (49, 50). Consistent with these roles, we detected an upregulation of deubiquitinase activity in the E9-high state (BH-adjusted *P* = 0.02, NES = 1.78).

**Fig. 4.**
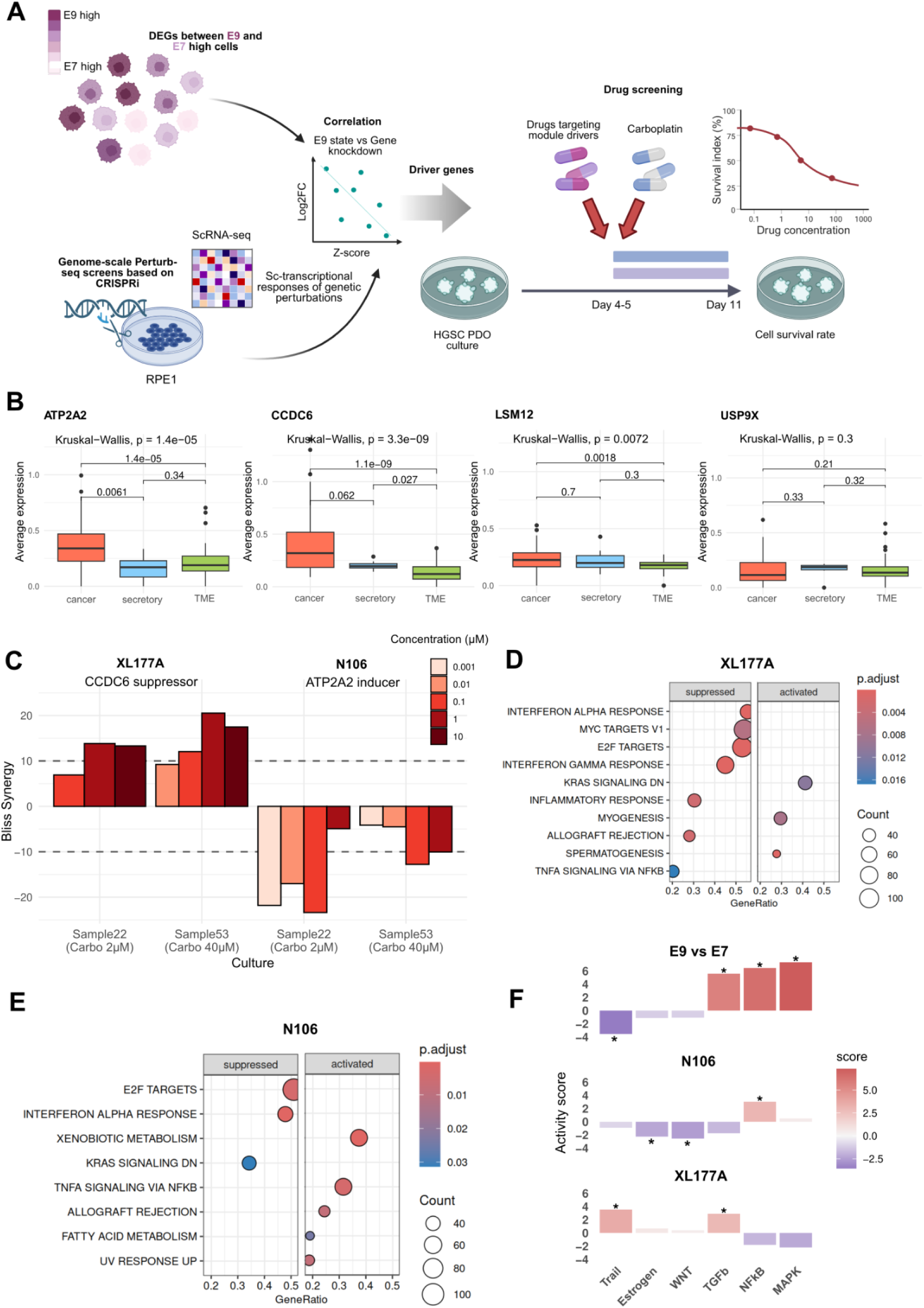
Small-molecule targeting of E9 state in PDO models. (**A**) Schematic representation of the identification of small-molecules targeting the modules utilizing expression profiles of genetically perturbed cells. LogFCs of gene module signature were compared with z-normalized expression of genes in perturbed data. The synergistic effects of module-targeting drugs combined with carboplatin were evaluated in PDO cultures. (**B**) Box plots showing the expression differences (Kruskal-Wallis test) of candidate target genes for state E9 in cancer and secretory epithelial cells and TME cells. Gene expressions were averaged per sample. (**C**) Bliss synergy scores for XL177A (*CCDC6* suppressor) and N106 (*ATP2A2* inducer) at carboplatin concentration 2 µM in sample22 and 40 µM in sample53 organoids. A synergy score greater than 10 was used as the threshold for synergistic interactions, while a score less than −10 indicated antagonistic interactions. (**D**) GSEA using hallmark gene set collection from MSigDB (with *P*.adjust < 0.05) showing gene sets activated or suppressed under XL177A treatment compared to untreated control. (**E**) GSEA using hallmark gene set collection from MSigDB (with *P*.adjust < 0.05) showing gene sets activated or suppressed under N106 treatment compared to untreated control. (**F**) Barplots showing the top three PROGENy pathways and their relative activity scores activated in E9 state compared to E7 state, and the effects of the drugs on these pathways. Pathways that remained significant after FDR correction of P-values are marked with an asterisk.

We performed drug testing in HGSC PDOs, derived from the same cohort as our scRNA-seq specimens, to assess the impact of *CCDC6*, *ATP2A2*, *LSM12* and *USPX9* on platinum response in a clinically relevant model. First, we analyzed the activity of the gene modules using previously published scRNA-seq data of HGSC organoid cultures (51). These cultures were derived from samples taken from five different patients: four from tumor tissue and three from ascites. We found that the E7 module was significantly more active in organoid cancer cells than in tumor tissue or ascites, while the E9 module is less active (P < 2.2e-16; Fig. S9A), consistent with the pseudotime analysis, as organoids should be biased towards early, stem-like cell states. In the drug screening, we targeted *CCDC6* via *USP7* suppression (52). For *ATP2A2*, we selected a SERCA2a activator, which is expected to enhance its function and thereby promote chemoresistance. A complete list of the tested drugs along with their target genes is provided in Table S5. Drug testing was performed on two PDOs (sample22 and sample53), with two technical replicates per condition. We examined the extent to which drug exposure affected cell survival when combined with chemotherapy. For the cell viability assays, organoid cultures were treated with the drugs both alone and in combination with carboplatin, and viability was assessed across four drug concentrations. Drug-response curves at the highest carboplatin concentration, along with dose-response matrices, are presented in Fig. S9B and Fig. S10. The results showed variability in treatment sensitivity across concentrations and organoid models. Using the Bliss independence model, we quantified synergy scores for drug combinations with carboplatin (Fig. S11A)(53). XL177A showed mainly synergistic interactions at 2µM carboplatin in sample22 and 40µM in sample53, while N106 induced mostly antagonistic interactions, especially in sample22 at the same carboplatin concentrations (Fig. 4C). Other drugs mostly had additive interactions across concentrations (Fig. S11B).

RNA-seq samples were collected after the sensitization period with candidate drugs, just before carboplatin treatment (see Methods). Principal component analysis (PCA) showed XL177A consistently induced a distinct gene expression profile compared to DMSO controls in both PDOs (Fig. S12A). The largest transcriptomic differences were due to organoid culture identity rather than treatment (Fig. S12B). Focusing on XL177A and N106, which displayed positive and negative synergy with carboplatin, respectively, we performed DGE analysis, accounting for organoid culture identity as a covariate. XL177A suppressed inflammatory signals, notably TNFα via NF-κB signaling (BH-adjusted P = 0.017, NES = −1.27; Fig. 4D), whereas N106, which is anticipated to enhance cancer cell resistance, activated this pathway (BH-adjusted P = 0.0034, NES = 1.75; Fig. 4E). Using a multivariate framework in which all PROGENy pathways were modeled jointly, we assessed the concordance between drug-induced pathway activation and our module profiles. NF-κB, one of the top pathways activated by module E9, was also found to be significantly activated upon N106 treatment (*P* = 8.53e-03). TNF-related apoptosis was the most strongly activated pathway in module E7, and it also emerged as the top pathway induced by XL177A treatment (*P* = 1.82e-03; Fig. 4F). These results suggest that the drugs elicit pathway-level responses that align with the transcriptional programs captured by the modules. Taken together, based on the gene modules we established, our analysis identified small molecules that impact cancer cell survival in the combination with a carboplatin treatment and target relevant resistance associated pathways.

## Discussion

Non-genetic heterogeneity is reflected in diverse transcriptomic states of cancer cells that manifest variable responses to drugs. The separation of these transcriptomic states in cancer cells is challenging as they are not as distinct as cell types. Transcriptomic states represent rather continuous signals and, therefore, clustering cells with similar transcriptional profiles, which produces distinct classification, is not the most optimal approach. To overcome this issue, cancer cell states can be characterized using co-expressed gene modules that are allowed to be expressed in combinations, highlighting the continuous range of possible states that can be achieved (23, 54), analogous to the Waddington landscape (55).

In our single-cell transcriptomic analysis, we explored the heterogeneity of cancer cells in HGSC using a co-expressed gene module approach. Unlike previous studies analysing cell states in HGSC (8, 10, 56, 57, 58), we incorporated transcriptomic data from both normal epithelial cells and components of the TME to dissect modules that are maintained, induced or lost during epithelial carcinogenesis. We identified two cell states derived from normal epithelia, reflecting contrasting phenotypes in OXPHOS and TNFα signaling, that were maintained through carcinogenesis, each having opposite effects on overall survival at the patient level. Notably, unlike the previous studies dissecting HGSC cell state heterogeneity, we further demonstrated that manipulating the identified states can influence chemotherapy responses *in vitro*.

TNFα signaling is known to have a dual role in cancer, promoting cell proliferation and tumorigenesis in some circumstances while inducing cell death in others (59). TNFα expression has been detected in 50% of HGSC patients, with a pathogenic role in inflammatory-induced ovarian carcinogenesis (60) and an upregulation of cytokines (61). Previous single-cell transcriptomic studies have identified EMT-enriched (12, 57) and stress-related (10) cell states that are significantly associated with poor prognosis in external cohorts. Our approach, which focused on epithelial-enriched modules, did not find patterns solely related to EMT. However, the NF-κB-TNFα pathway scores were highly aligned with those of TGF-β signaling that is a major driver of EMT in both the cancer hallmark and progeny pathway analyses. Furthermore, *TNF* was one of the markers of the stress-associated cancer cell state we identified earlier to be induced by neoadjuvant chemotherapy, promoting a vicious signaling cycle between stress-associated cancer cells and inflammatory cancer-associated fibroblasts (10). Jointly, these results suggest that the poor prognosis state with increased TNFα-NF-κB signaling identified is reflecting the key features of previously identified resistant states related to stress (10) and EMT (12, 57), supporting the robustness of our approach.

Our results associate the pseudotime-early module E7 with increased OXPHOS associated gene expression, HRD genotype and a favourable prognosis. OXPHOS has been connected to HRD in both patient (62) and functional levels (63), to better prognosis in both animal models and a clinical cohort (63), with opposing results in the cell line context (64). Epithelial and cancer stem cells have been shown to prefer OXPHOS metabolism (65) and *BRCA1* mutations enhanced stem-like properties in breast epithelial cells (66). Although early, stem-like states are classically linked to resistance and metastasis, recent colon cancer studies suggest that metastasis instead involves reversal of stemness (67), leading to aberrant differentiation programs in resistant metastases (68). Consistently, the pseudotime-late, poor-prognosis state we identified showed enrichment for epidermal differentiation genes. Combined, these earlier studies and our data support closely intertwined links between pseudotime-early cell state, HRD, OXPHOS and a favourable prognosis, with mutual exclusivity to TNFα signaling, in line with a previous study (69).

Although our analysis focused on the contrasting patterns that were maintained through carcinogenesis, we also found differences in their representation between normal and cancer tissues. For example, antigen processing and presentation were suppressed from normal to cancer in the pseudotime-late E9 state. This may reflect the highly antigen presenting, senescent phenotypes of late-pseudotime cells in normal epithelia that are suppressed in cancer to enable immune evasion (70). Uniquely, our cohort included a normal fallopian tube specimen from a patient with a germ-line *TP53* mutation. Despite low *TP53* expression, these cells resembled normal epithelia rather than cancer across the co-expression modules, also maintaining normal-like antigen presentation patterns in the pseudotime-late state. This is in line with proteomic and morphologic analyses of P53 signature cells that precede STICs in HGSC carcinogenesis (71, 72). Hence, even though *TP53* is the initial genetic driver of HGSC carcinogenesis, further aberrations are required for the emergence of cancerous phenotypes, including immune evasion.

As we only have associative rather than causal evidence for the individual module genes’ role in resistance, we used public CRISPR-perturbed expression profiles to unbiasedly identify potential drivers that allow functional targeting of these states. *CCDC6* was the top predicted driver of the poor-prognosis state and showed the strongest negative link to the good-prognosis state. *CCDC6* maintains genomic integrity by regulating protein phosphatase 4 complex activity on the DNA damage indicator γH2AX, and its loss leads to PARPi sensitivity and HRD of HGSOC cells (73). In line with the role of *CCDC6* driving HRD, and predicted opposition of the good prognosis E7 state, its targeting the small molecule XL177A showed synergy with carboplatin treatment in PDOs. *CCDC6* loss may further influence NF-κB-TNFα signaling via the protein phosphatase 4 complex (74), in line with NF-κB-TNFα suppression following XL177A treatment in the PDOs. Our experiments further confirmed that as predicted, the SERCA activator N106 induces carboplatin resistance. SERCA controls ER stress (75) that is linked to poor outcomes in ovarian cancer (76). While NF-κB inhibition was previously shown to reduce SERCA activity and increase ER stress (77), we found that SERCA activation induced NF-κB signaling, suggesting a feedback loop. Overall, our approach effectively identified targetable cell state drivers that influence key resistance pathways and chemosensitivity in PDOs.

In conclusion, our module-based profiling of HGSC elucidates a prognostic role of conserved epithelial cell states shared between normal FTE and cancer. We identified that TNFα/NF-κB associated late state predicts poor clinical outcome in HGSC patients and can be targeted to modulate the chemosensitivity of PDOs. Our approach leverages single-cell transcriptomic data from HGSC and its cell of origin in a novel and integrative manner to gain new, actionable insights into cancer biology.

### Limitations of the study

The limitations of this study primarily relate to the challenges of harmonizing single-cell data from heterogeneous cancer samples, the constraints of available drug prediction databases, and the resource-intense nature of generating and maintaining PDO models. HGSC cells exhibit substantial inter-patient heterogeneity, and the number of captured cancer cells varied markedly across samples. The variability poses a challenge for assessing gene co-expression patterns even though they are derived from pseudobulk-level DEGs, as sample-specific effects may confound the identification of broadly conserved transcriptional programs. For module driver gene identification, we utilized publicly available Perturb-seq data from RPE1 cell line. Although this cell line has an epithelial origin, it does not fully recapitulate the gene expression dynamics of HGSC cells either in tumors of the PDOs used for drug testing, limiting the direct translatability of perturbation effects. To evaluate therapeutic relevance, we targeted gene modules in PDO models, which are considered more predictive of patient-specific responses than conventional cell lines. Only two distinct organoid models were available for this study, which limits scalability and reduces the generalizability of the findings across broader patient populations. In addition, RNA-seq was performed at a single time point after drug treatment prior to carboplatin exposure, which restricts our ability to evaluate the timing and duration of the observed sensitization effect.

## Methods

### Study cohort

All patients, who contributed HGSC or normal FTE samples for the study, provided written informed consent. The Ethics Committee of the Hospital District of Southwest Finland approved the study and the use of all clinical materials under decision number EMTK: 145/1801/2015. HGSC patients were treated at the Turku University Hospital in Finland and tumor samples were collected from the patients as part of the DECIDER study (NCT04846933). Our data cohort comprises 16 previously unpublished scRNA-seq samples and 43 previously published samples reported in (10, 51, 78, 79, 80). Detailed publication information is provided in Table S1.

### scRNA-seq sample preparation

HGSC tumor samples (n = 52) were collected from the patients in laparoscopy or primary debulking surgery (PDS) depending on the treatment strategy. Non-cancerous fallopian tube or fimbria tissue samples (n = 7) were collected from post-menopausal women at the time of other gynecological operations. Instantly after surgery, the samples were incubated overnight in a mixture of collagenase and hyaluronidase to obtain single-cell suspensions (Department of Pathology, University of Turku). Single-cell suspensions were frozen in STEM-CELLBANKER DMSO-FREE solution and thawed in culture medium immediately prior to processing for scRNA-seq. The viability of the frozen single-cell suspensions ranged from 26.4 to 94.9% with a median of 83.1% after thawing. The standard Chromium Single Cell 3’ Reagent Kit v.2.0 (10x Genomics) was used for scRNA-seq library construction and they were sequenced on Illumina HiSeq4000 (Jussi Taipale Lab, Karolinska Institute, Sweden), HiSeq2500 and NovaSeq6000 sequencing instruments (Sequencing Unit of the Institute for Molecular Medicine Finland, Finland).

### Preprocessing scRNA-seq data

Demultiplexing, alignment, barcode filtering and gene expression quantification based on UMIs were performed by the Cell Ranger software (version 3.1.0). The reference index was constructed based on the GRCh38.d1.vd1 genome annotated with GENCODE v25. Seurat v4 R toolkit was used to preprocess filtered feature-barcode matrices (81). We filtered out cells having more than 20% of UMI counts originating from mitochondrial genes. To remove technical variability, we used sctransform as a normalization method. PCA was conducted by computing 50 PCs, and UMAP projection was generated with 30 dimensions. Major cell types were identified based on canonical markers: epithelial cancer cells (*MUC16, PAX8, WFDC2*), epithelial ciliated cells (*FOXJ1, PIFO*), epithelial secretory cells (*OVGP1, PAX8*), stromal cells (*COL1A2, DCN, FGFR1, PECAM1, VIM*) and immune cells (*CD14, CD3D, CD79B, FCER1G, HLA-DRA, MZB1, NKG7, PTPRC, HBB*). Cells showing any expression of stromal markers *DCN* or *LUM*, or immune markers *CD3D, CD79B, FCER1G, HBB* or *PTPRC*, in addition to epithelial markers *MUC16* and *PAX8* were removed as doublets. Shared nearest neighbor (SNN) modularity optimization based clustering algorithm with resolution 0.5 was used to identify clusters and cell types were assigned to the clusters according to the markers. Ciliated cells were excluded from further analysis. The quality metrics were calculated for each cell, and QC threshold was determined for each cell type based on its bimodal form in the 2D density plot. Cell type-specific thresholds for log2nCount were 11 for epithelial cells, 11 for stromal cells and 10 for immune cells. In setting the threshold for epithelial, it was taken into account that the control samples had, overall, lower counts compared to the cancer samples. After cell type-specific cell filtering, the normalization, dimensionality reduction, clustering and cell type assignment steps were repeated, resulting in a total of 131 225 cells, including 30 236 malignant epithelial, 1 251 secretory FTE, 80 764 immune, and 18 974 stromal cells.

### Differential gene expression analysis and genes for the gene modules

We used edgeR - limma workflow (v.4.2.0) to perform pseudobulk differential gene expression (DGE) to ensure also that lowly expressed genes are detected (82). Pseudobulk expression was calculated only for samples having more than 50 cells per cell type and therefore 9 tumor samples and 2 FT samples were excluded from this analysis. In addition, an external FT scRNA-seq data set containing 3 614 secretory cells from 7 patients was utilized to get a bigger secretory cell count to the DGE comparison (19). We followed standard limma-voom workflow including calculation of normalization factors, voom transformation, calculation of variance weights, fitting linear model and ranking genes based on an empirical Bayes method. DEGs were calculated in three comparisons: between cancer and secretory FTE cells, between cancer cells and cells from the TME, and between secretory FTE cells and cells from the TME. DGE results were further filtered to ensure that genes and their expression have a functional role and to represent biological differences between the cell groups. Genes with an adjusted *P*-value greater than 0.05 were excluded. Genes had to be expressed in at least 10% of cells in the cell group and two-fold compared to the comparison group. Genes had to be protein-coding but not ribosomal or mitochondrial genes. Additionally, cancer- and normal FTE-specific genes had to be overexpressed in either cancer or normal FTE cells compared to cells from TME. Alongside cancer- and normal FTE-specific gene groups, we identified a shared epithelial gene group consisting of genes overexpressed in both cancer and normal FTE when compared separately to cells from TME. Based on these criteria, we identified 229 cancer-specific, 257 normal FTE-specific and 598 shared epithelial genes.

### Identification of co-expressed gene modules

To infer co-expression for three identified gene groups, we employed a statistical method CS-CORE with R package implementation (31). We followed their vignette to implement an iteratively re-weighted least squares approach for estimating correlations between expression levels controlling confounding effects extracting co-expressed gene modules. Co-expressions that were not statistically significant using the Benjamini-Hochberg procedure were set to zero. Co-expression for cancer-specific genes was calculated using all cancer cells, while co-expression of normal FTE-specific genes was assessed in secretory FTE cells, including data from external scRNA-seq source (19). Co-expression for epithelial genes was initially calculated separately for cancer and normal FTE cells, and then averaged to mitigate the impact of cell count imbalance between these two cell groups. We applied traditional WGCNA workflow to extract co-expressed gene modules from the thresholded co-expression matrices (32). First, the adjacency matrix was computed from co-expression matrix, and then it was transformed into topological overlap matrix (TOM), which does not consider each pair of genes in isolation but also captures how similarly they are connected to other genes in the network. The TOM was converted into a dissimilarity matrix for hierarchical clustering with the average linkage method. The dynamic tree cut algorithm was used to cut the dendrogram into gene modules (83).

### Cellular activities of gene modules and pathway activity inference

We calculated gene module activities in single cells using the UCell scoring method (R package v2.8.0) with default parameters (33). UCell employs a rank-based approach to compute single-cell gene expression scores from a gene set, which we found to effectively quantify module activities across different samples and experimental data. Hallmark gene sets from the MSigDB were utilized to examine biological functions of the gene modules. Cells were scored with UCell algorithm using hallmark gene sets, and Spearman’s correlations were calculated between the gene modules and hallmark gene sets separately in cancer and normal FTE cells (34). Correlations with adjusted P > 0.05 were set to zero. To reduce sample-specific impact, a subset of cancer and normal FTE cells was used for comparison, where cells were selected per sample based on the median cell count (236 for cancer cells and 151 for normal FTE cells).

We used PROGENy (R package v1.26.0) implementation to infer the activities of 14 signaling pathways using consensus gene signatures obtained from perturbation experiments when we studied differences in E7 and E9 modules in cancer cells (39). Spearman’s correlations were calculated between the gene module and pathway activity scores at a single-cell level, and adjusted P values were calculated for the correlations. We calculated pathway correlations with the gene modules also in the external scRNA-seq data set by Vázquez-García et al. using all 211 624 HGSC cells from 41 patients (8).

Gene set enrichment analysis (GSEA) was implemented through clusterProfiler v4.12.0 R package to study enrichment of GO terms in several comparisons related to the gene modules (84). Gene lists were ranked based on log fold change (logFC) values obtained from the Seurat *FindMarkers* function or the limma *toptable* function depending on whether single-cell or bulk data were used. In the gseGO function, 10000 permutations were performed for robust statistical assessment, and gene sets containing between 3 and 800 genes were included. Enrichplot v1.24.0 was used to make dot plots from enrichment results.

### RNA velocity analysis

Alignment and quantification of the spliced and unspliced mRNA for each gene was performed with velocyto v0.17.17 in Python v3.8 (85). To mask repetitive elements, a repeat annotation file downloaded from UCSC genome browser was used. The velocyto read counting pipeline was run through *velocyto run* command. To estimate transcriptional states and infer velocity of RNA changes for individual cells, we used the dynamical model from scVelo v0.2.3, which incorporates information from both spliced and unspliced mRNAs (37). Cells were ordered along a pseudo-temporal trajectory using root and terminal cells identified with CellRank v1.3.1 (36). To improve root cell identification, we used a symmetric graph in a velocity graph construction with the function *scv.tl.velocity_graph*(mode_neihbors=’connectivities’). The identification of initial and terminal states as well as latent time scoring were performed separately for those samples having more than 100 cells. The RNA velocity analysis was also performed for the control samples including ciliated cells. Spearman’s correlation was used to calculate association between latent time and gene module scores on a sample-wise basis. Only the correlations having adjusted P < 0.05 were used when averaged correlations between latent time and gene modules were computed across the samples.

### Bulk TCGA RNA-seq data and HRD signature

Deconvoluted bulk TCGA RNA-seq data from primary tumors of 258 HGSC patients (grade: G3 and G4) with available OS data was used as a cohort to study the prognostic impact of the gene modules (10, 35). For survival analysis, TCGA patients were divided into two groups based on an optimized module score threshold using the max.stat v0.7.25 R package (86). TCGA patients were classified into HRD and HRP groups using ovaHRDscar genomic classifier (87). BRCAness transcriptional signature from COV362 ovarian cancer cell study was used to validate the HRD association of module E7 (11). The BRCA-mutated (BRCAmut) gene set, consisting of 838 genes, and the BRCA wild-type (BRCAwt) gene set, consisting of 249 genes, were constructed based on logFC threshold zero and, they were compared to the module signature using GSEA. Copy-number alterations inferred by GISTIC for TCGA cohort were downloaded from cBioPortal (88, 89, 90). Copy number alteration profiles were found for 248 patients out of 258. Only frequently altered genes in HGSC reported previously (44), that were amplified in at least 10 TCGA patients, were included in the module activity analysis.

### Module active cells and the construction of module signatures

We determined as module active cells those top 25% of cells having the highest UCell score. We also tested thresholds 10%, 25% with cells subsetted per sample by mean cell count, and 50% for module active cells and identified differentially expressed genes between these cells and all other cells. For this analysis to get module specific signatures, we used the Seurat function *Findmarkers*(test.use = ‘LR’, latent.vars=c(‘sample’, ‘S.Score’, ‘G2M.Score’), logfc.threshold=0). The same method was used to identify genes differentially expressed between E7- and E9-module active cells where only cells categorized either E7 active or E9 active were used. When E9-module active cancer cells were compared to normal FTE cells, MAST-test was used in the DGE analysis as it was able to regress out sample-specificity with separate test groups. In this comparison, 234 DEGs found also in pseudobulk comparison between cancer and normal FTE were filtered out before GSEA to reduce signal originating from differences between these cell types. All the DGE results were filtered to retain only genes with an adjusted *P* value < 0.05.

### Inference of CNV profiles and subclones

The CNV profiles and subclones were inferred using the InferCNV (v1.7.1 R package) (45). In the *run* function, we used the following parameters: cutoff = 0.1, cluster_by_groups = T, analysis_mode = ‘subclusters’, denoise = T, leiden_resolution = 0.5, HMM = T, HMM_report_by = ‘cell’. The subclone identity was calculated for each cell. We used immune and stromal cells from each patient as a reference for inferring CNVs. Only patients having more than one subclone and subclone groups having more than 10 cells were used in the Chi-square test. Altogether, a total of 10 615 cells with either module E7 or E9 active and with available subclone information were used in the analysis. A Chi-square test was performed both within individual patients and across all patients to assess the distribution of the variables.

### Module targeting based on Perturb-seq screens

To identify small-molecules to target gene modules, we used an *in silico* approach and utilized genome-scale Perturb-seq screens (46). We used retinal pigment epithelial (RPE1) cell line data consisting of single-cell transcriptional responses for 2394 knockdown genes. We calculated Spearman’s correlations between z-scores of perturbed genes and logFCs of module signatures. In the comparison, 7190 genes were used with the E7 module and 7229 genes with the E9 module. The correlation coefficients having adjusted P value > 0.05 were filtered out. Genes with the most negative correlations with the gene modules were considered module drivers as Perturb-seq screens captured transcriptional responses of gene silencing. Given that the E7 module demonstrated a positive effect on survival and our objective is to promote this cellular state, we prioritized genes exhibiting the strongest positive correlations. These gene candidates were further required to be differentially expressed in module-high cells compared to the other cells.

### Organoid drug screening

Organoids used in the experiments were established, genomically validated to represent original tumor samples and contain only cancer cells and cultured as previously described (51). For drug tests, organoid cultures were trypsinized to obtain the suspension of organoid fragments. The cells were resuspended in the Cultrex BME-2 matrix [#3533-010-02, Biotechne], dispensed to 384-well Ultra-Low Attachment microplates (#4588, Corning) at approximately 2 x 10^3^ cells per well in 10 µl of the matrix, and covered with 40 µl of growth medium containing 5 µM ROCK inhibitor (#HY-10071, MedChemExpress) to facilitate the organoids formation. After 2–3 days, the medium was exchanged to ROCK inhibitor-free growth medium. After additional 2-3 days of culture, the medium was exchanged and the tested compounds (10-fold dilution series of four concentrations, ranging from 10 nM to 10 μM), carboplatin (organoid line-specific, see figures), vehicle (DMSO), or positive control compounds (10 μM bortezomib) were transferred to the wells using Echo 550 acoustic dispenser (Labcyte). The organoids were incubated with drugs for a total of 7 days (with medium exchange and drug replenishment after 4 days) in the cell incubator at 37 °C. At the endpoint/readout, 25 μL of the culture medium was exchanged to staining solution, containing CellTox Green reagent (#G8743, Promega, final concentration 1X) and Hoechst 33342 (#B2261, Sigma, final concentration 5 μg/mL). Plates were incubated for 2h at room temperature and imaged in an ImageXpress Micro Confocal automated fluorescence microscope (Molecular Devices). Images were analyzed using and cytotoxicity indices were calculated using MetaXpress software, as described previously (51).

Drug combination effects were calculated using the Bliss independence model [Ianevski]. Cell viability results were normalized prior to Bliss score calculation to minimize data variability and to prevent plate-dependent spurious synergy effects. Synergy scores greater than 10 were interpreted as evidence of synergistic interactions, while scores below −10 indicated antagonism.

### RNA-seq on organoids

For RNA-seq, confluent organoids growing in 6-well plates (#140685, Thermo Fisher Scientific,) were harvested and passaged as previously described (51). In short, hydrogel domes were mechanically and enzymatically digested in 2 mL of TrypLE Express Enzyme (#12604013, Thermo Fisher Scientific,) and incubated for 15 min in the cell incubator at 37°C. The cell digests were collected in a conical tube and each well was rinsed with 1 mL of PBS. After centrifugation at 300 x g for 5 min, the supernatant was carefully removed. To seed the organoids, the cell pellets were resuspended in Cultrex BME-2 matrix [#3533-010-02, Biotechne], with the protein concentration adjusted to 8 mg/mL with ice-cold PBS. The seeding density followed the particular split ratio of each organoid culture and a minimum of 12 wells of a 6-well plate were seeded, with each well containing 10 domes of 20 μL. After the seeding, organoids were maintained in culture medium (M1 or M2) supplemented with Rho-associated protein kinase inhibitor (#HY-10071, MedChemExpress) at a final concentration of 5 µM. On day 4, the media was exchanged to ROCK-free growth medium containing the test compounds. The compounds had been reconstituted in DMSO and were used for the treatments at the following final concentrations: XL177A (1 μM), trans-ned-19 (10 μM), N106 (1 μM), EOAI3402143 (1 μM), and FT709 (10 μM). Control wells (drug-free) were treated with DMSO (0.1% v/v). Overall, the drug treatments lasted for 48 hours, and each condition conducted in technical duplicates. On day 6, organoids were harvested as previously described, keeping the technical duplicates independent. After centrifugation, pellets were harvested in 400 μL of 1X DNA/RNA ShieldTM reagent (#R1200, Zymo Research), transferred into a microcentrifuge tube, and stored at −20°C. RNA was isolated following the “Total RNA isolation” protocol from the Quick-RNA Miniprep Plus Kit (#R1058, Zymo Research). After isolation, the RNA purity and concentration were assessed with a spectrophotometer (NanoDrop2000, Thermo Scientific). Sequencing of the messenger RNA (mRNA) was conducted by Novogene Europe (Planegg, Germany) after quality control. Library preparation included selection of mRNA from the total RNA by poly-A enrichment. The sequencing was done with the NovaSeq X Plus Series platform from Illumina using a paired-end sequencing kit of 300 cycles (PE150) resulting in 20 million read pairs per sample.

### RNA-seq processing

Paired-end RNA reads were quality controlled with FastQC (v0.12.1) [91]. Adapters, poly-A tails and bases with a Phred score of less than 20 were removed using Trim Galore! (v0.6.10) with cutadapt (v4.9) (92), removing reads with less than 20bp left after trimming. Trimmed reads were aligned with STAR (2.7.11b) (93) to the human reference genome GRCh38.p14 primary assembly. Transcripts were quantified using Salmon (v1.10.3) (94) in alignment-based mode and summarised to gene transcripts per million (TPM) using the Ensembl (Release 113) (95) gene database.

The limma workflow described earlier was used to find genes differentially expressed between drug treated and control organoid samples. Organoid ID was included as a covariate in the design matrix to account for inter-organoid variability. PROGENy pathway activity inference for the organoid RNA-seq samples was performed using decoupleR v2.10.10 and the multivariate linear model method.

### Statistical analyses

Wilcoxon rank-sum test was used for pairwise comparison and Kruskal-Wallis test for the comparison of multiple groups. They were produced using the *stat_compare_means* function from the ggpubr v0.6.0 R package for the figures. Shapiro-Wilk test from stats v 4.4.1 R package was used to test normality of data.

Survival analysis was computed using survival 3.7.0 and survminer 0.4.9 R packages. Kaplan-Meier curves were generated to analyze overall survival (OS). OS was defined as the time from diagnosis of HGSC to death or the last follow-up. Multivariate Cox proportional-hazard model was used to describe how gene module activity, HRD status and patient’s age jointly impact on OS.

## Data availability

The unpublished raw scRNA-seq data will be deposited in the European Genome-phenome Archive (EGA), and the processed data will be submitted to the Gene Expression Omnibus (GEO) upon publication.

## Code availability

The scripts used for scRNA-seq analyses will be made publicly available via our GitHub repository (https://github.com/XXX) upon publication. The Python and R packages employed in this study are listed in the Methods section and are publicly accessible online.

## Author contributions

Conceptualization: A.P. and A.V. Methodology: A.P. and A.V. Investigation: A.P., A.V., L.G-M., W.S, D.F-A., Formal analysis: A.P., D.F-A. and M.M.F. Validation: L.G-M., D.F-A. and W.S. Resources: J.H., K.W. and A.V. Data curation: M.M.F., E.P.E., J.H. and A.V. Visualization: A.P. Supervision: A.V. Project administration: A.V. and K.W. Funding acquisition: A.V. and K.W. Writing—original draft: A.P. and A.V. Writing—review & editing: A.P., A.V., L.G-M. and E.P.E.

## Acknowledgments

Data was acquired in the DECIDER project, which was funded by the European Union’s Horizon 2020 research and innovation programme under grant agreement No 965193. This study was co-funded by the European Union (ERC STRONGER 101125261). Views and opinions expressed are however those of the author(s) only and do not necessarily reflect those of the European Union or the European Research Council. Neither the European Union nor the granting authority can be held responsible for them.

Funding was also received from the Sigrid Jusélius Foundation, Research Council of Finland (Number 344697 for Era PerMed JTC2020 project PARIS (Precision drugs Against Resistance In Subpopulations, and Number 351196), Cancer Foundation Finland, the Finnish Cancer Institute (K. Albin Johansson Cancer Research Fellow position for AV), and HORIZON-MSCA-2021-PF, European Commission (M.M.F.: 101067835). The authors thank the CSC – IT Center for Science for computational resources.

The results published here are in part based on data generated by the TCGA Research Network: https://www.cancer.gov/tcga.

---

**Fig. S1.**
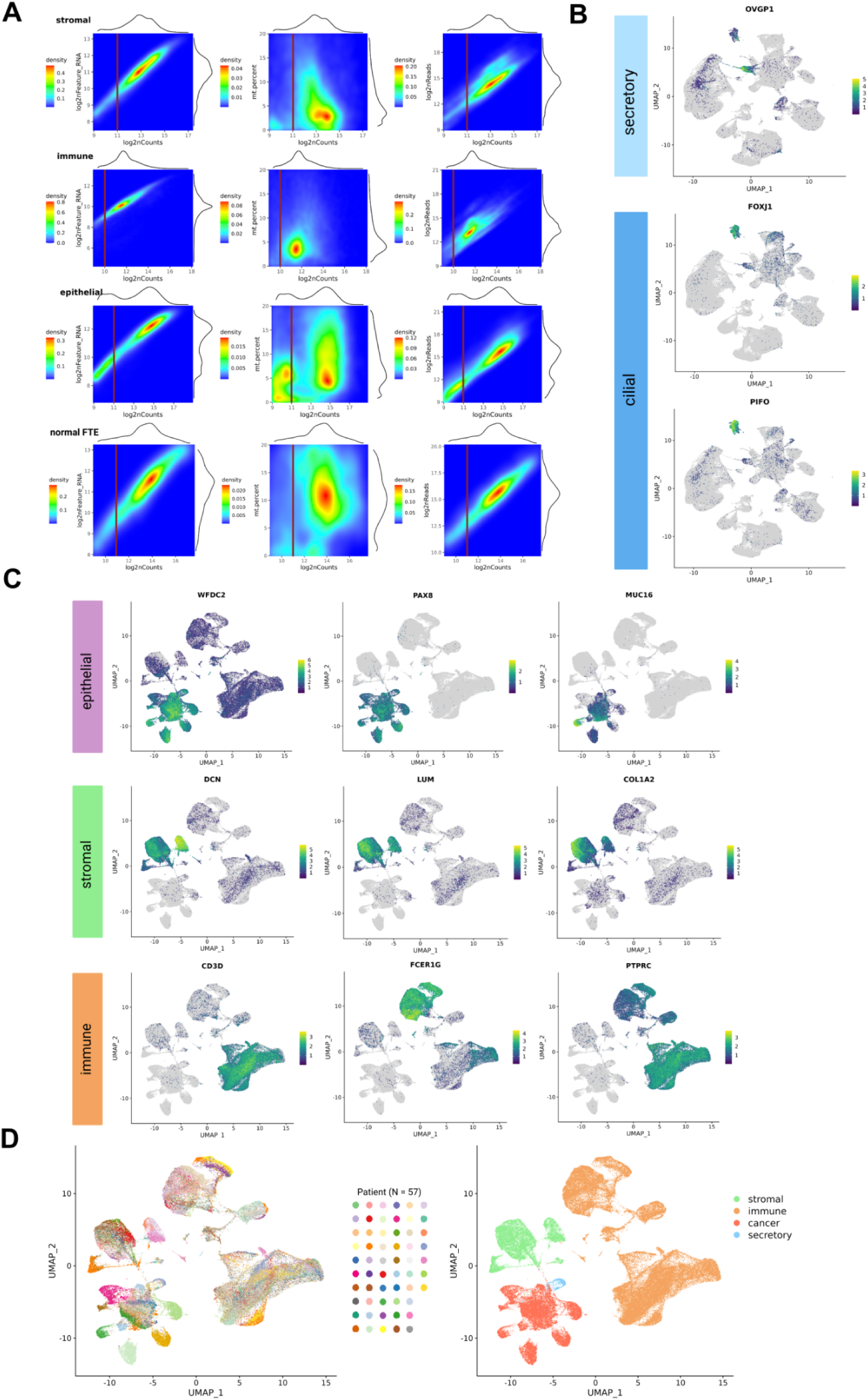
Quality control for scRNA-seq data and identification of normal epithelial, cancer, stromal and immune cells. (**A**) Density plots showing quality control (QC) metrics (log2nCounts, log2nFeature, percentage of mitochondrial derived counts (mt.percent)) and the cutoffs for them in stromal, immune, epithelial and normal FTE cells. (**B**) UMAPs showing the expression of FTE cell markers in remaining cells after quality control. Yellow indicates higher expression of a gene while blue indicates lower expression. Gene is not expressed in the cells colored by gray. (**C**) UMAP of 131 225 cells after filtering cilial type of FTE cells and double-marker positive cells showing the expression of canonical markers of cancer epithelial, stromal and immune cells. (**D**) UMAPs of cells colored by patient (left) and by cell type (right).

**Fig. S2.**
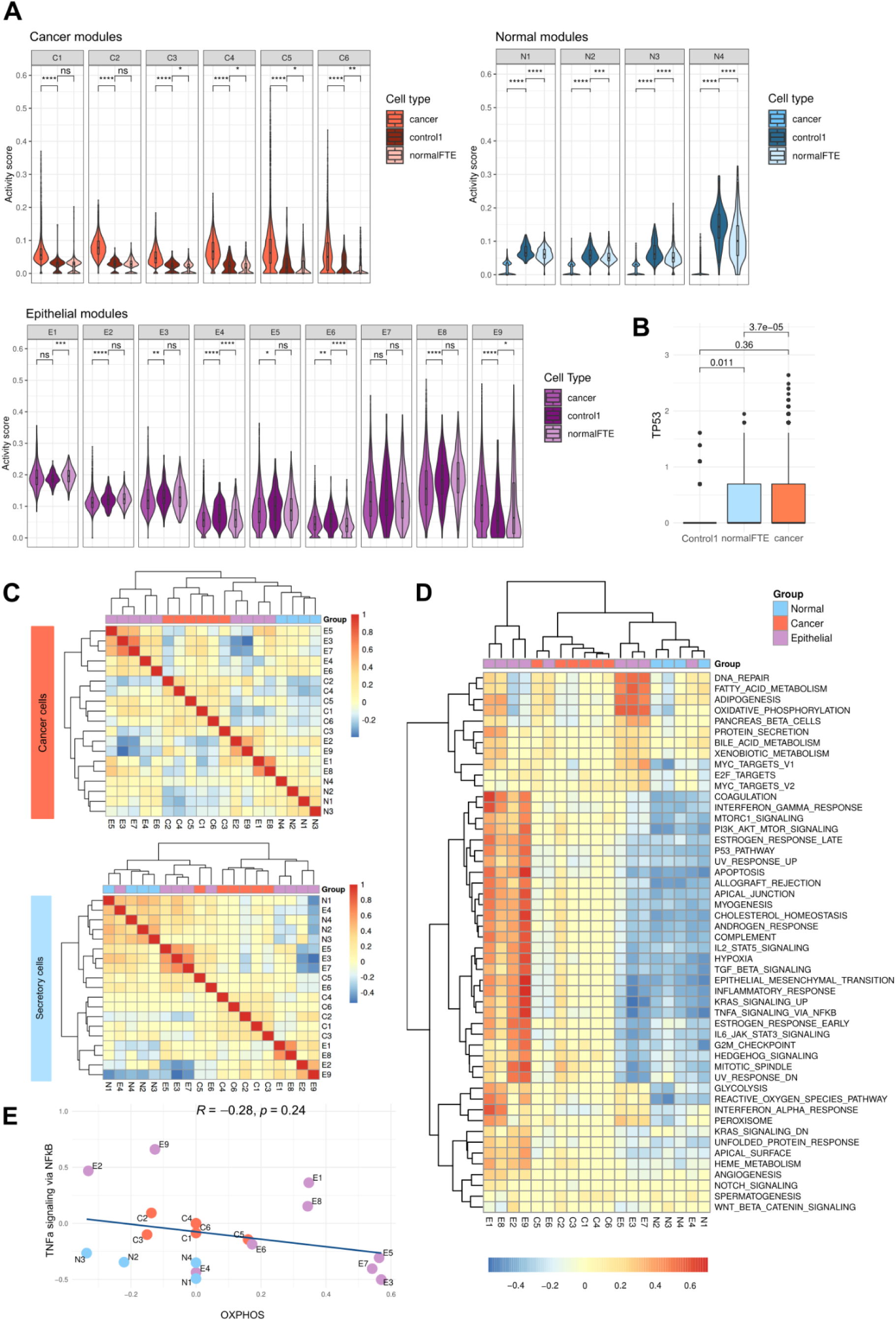
Gene module similarities and the characteristics of gene modules in normal epithelial cells. (**A**) Violin plots showing module activities in p53-mutated normal sample control1 (n = 151) compared to cancer and other normal FTE. Pairwise comparisons were performed using the Wilcoxon test. (**B**) Box plot showing *TP53* expression in control1 compared to normal FTE and cancer cells. (**C**) Correlation heatmap of gene modules calculated at a single-cell level using UCell scores in cancer cells (n = 8 786) (left) and secretory FTE cells (n = 712) (right). (**D**) Correlation heatmap between gene modules and hallmark gene set collection from MSigDB calculated at a single-cell level of secretory FTE cells (n = 712) using UCell for the calculation of signature activity. (**E**) Correlation plot of TNFα signaling via NF-κB and OXPHOS across all the gene modules (n = 19) in secretory FTE cells, showing no relationship. Data points are colored by module group.

**Fig. S3.**
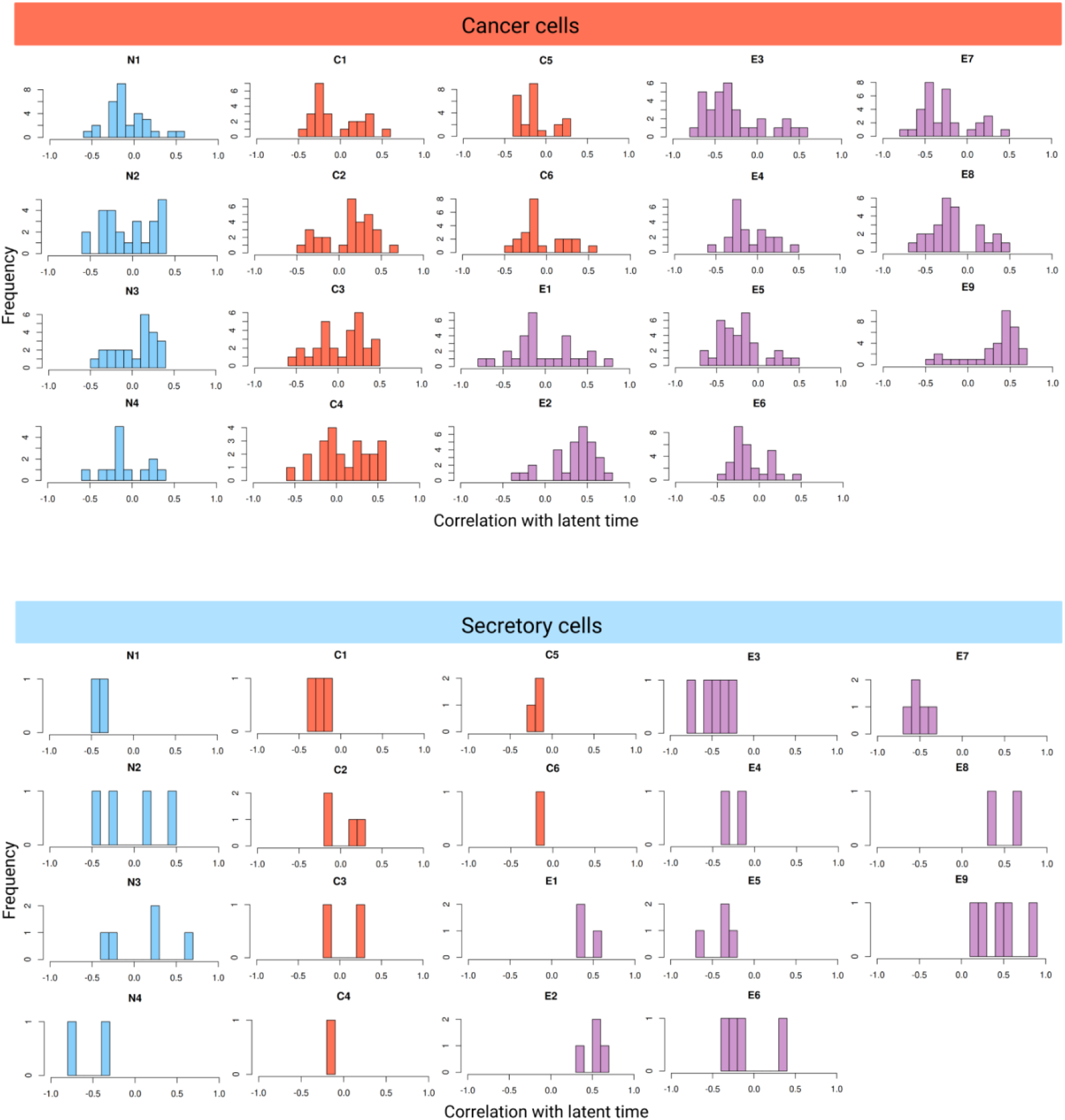
Latent time distributions of gene modules in cancer and secretory cells. Histograms showing distributions of sample specific correlations with a latent time for the gene modules in tumor samples (above) and in FTE samples (below). Only correlation coefficients having *P* < 0.05 after FDR-correction are shown in histograms.

**Fig. S4.**
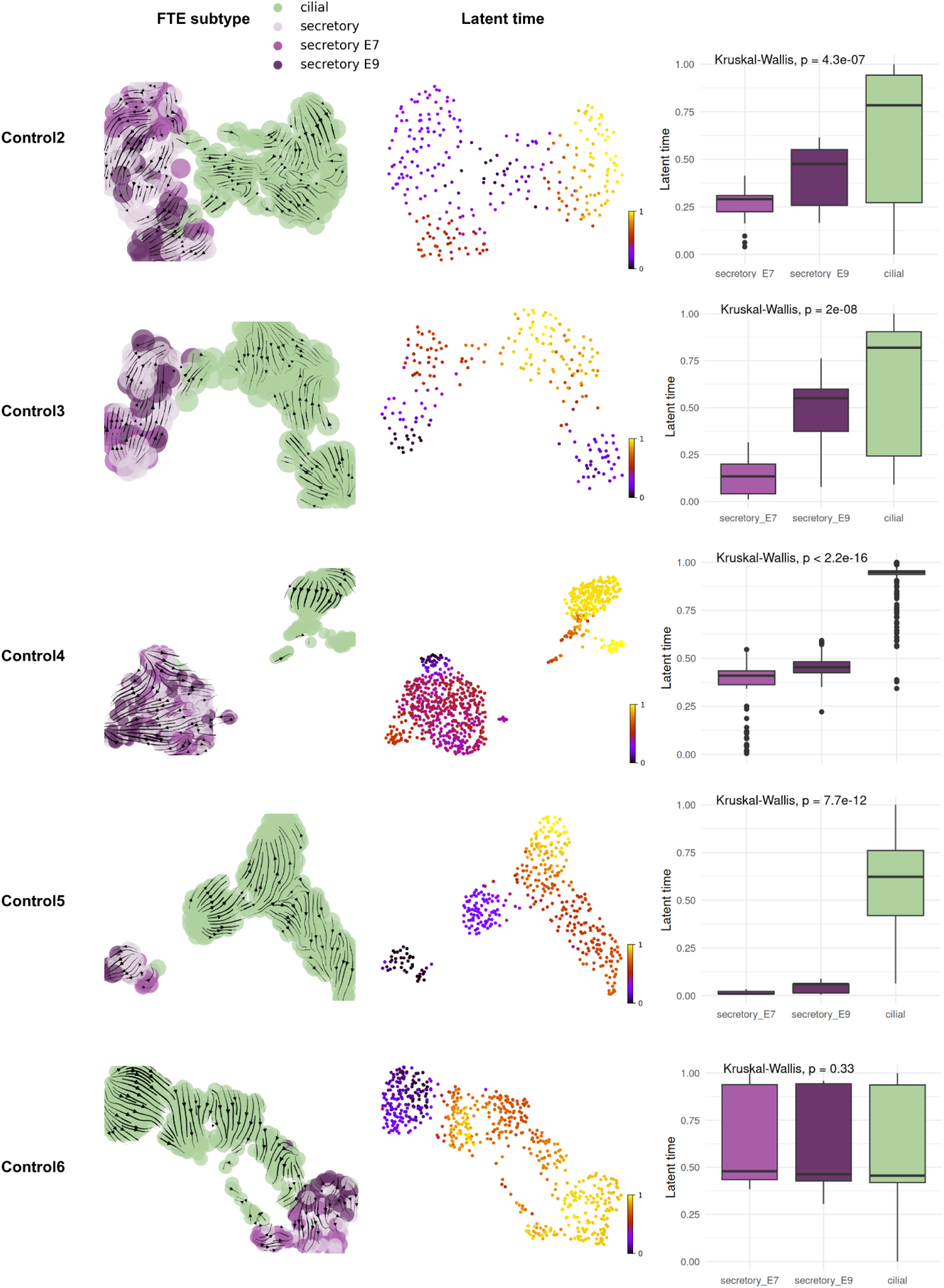
Latent time analysis of cells from control samples indicates that the ciliated state follows the secretory state in FTE cells. UMAPs of normal FTE secretory and ciliated cells in five control samples. Cells are color-coded by FTE subtype, and the top 25% of cells with the highest activity of either E7 or E9 state are also indicated (left), alongside their latent time scores (right). Velocity vectors and latent time are calculated with the CellRank algorithm. Box plots show the differences in latent time in secretory cells with the highest activity in E7 or E9, and ciliated cells.

**Fig. S5.**
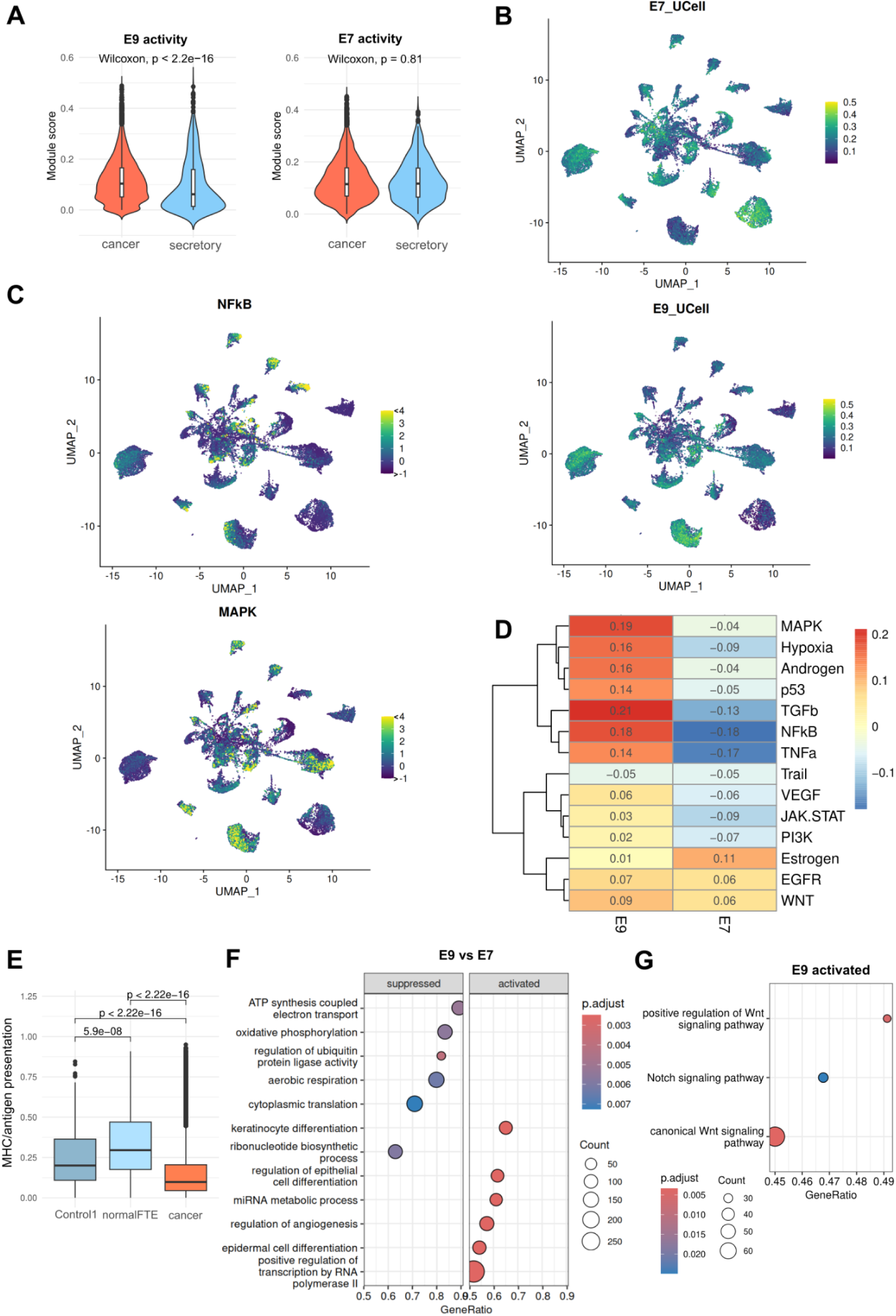
Biology of the E7 and E9 states. (**A**) Violinplot comparisons of E9 and E7 activities in cancer and secretory epithelial cells (Wilcoxon rank-sum test; P < 2.2e-16 and 0.81, respectively). (**B**) UMAPs of cancer and secretory epithelial cells colored by the E7 (above) and E9 (below) module activities calculated with UCell. (**C**) UMAPs colored by the level of NF-κB signaling pathway (above) and MAPK signaling pathway (below) activities calculated by PROGENy. (**D**) Correlation heatmaps of pathway activities, inferred using PROGENy, were generated for E7 and E9 in HGSOC cells (n = 211624 from 41 patients) from the external dataset by Vázquez-García et al. (**E**) Box plot showing MHC protein complex/antigen presentation signature activity calculated from normal-enriched E9 DEGs contributing these pathways (*CD74, HLA-C, HLA-DMA, HLA-DPA1, HLA-DPB1, HLA-DQA1, HLA-DQB1, HLA-DRA, HLA-DRB1*) in control1 compared to normalFTE and cancer cells (**F**) GSEA scores for GO terms (with Bonferroni-adjusted *P* < 0.05) were calculated between E7 and E9 modules using 25% of cancer cells per sample showing the highest activity of the module. (**G**) GSEA scores for Notch and Wnt signaling pathways activated in the E9-high state (from top to bottom BH-adjusted *P* = 0.0052, NES = 1.92; BH-adjusted *P* = 0.025, NES = 1.68; BH-adjusted *P* = 0.0034, NES = 1.84).

**Fig. S6.**
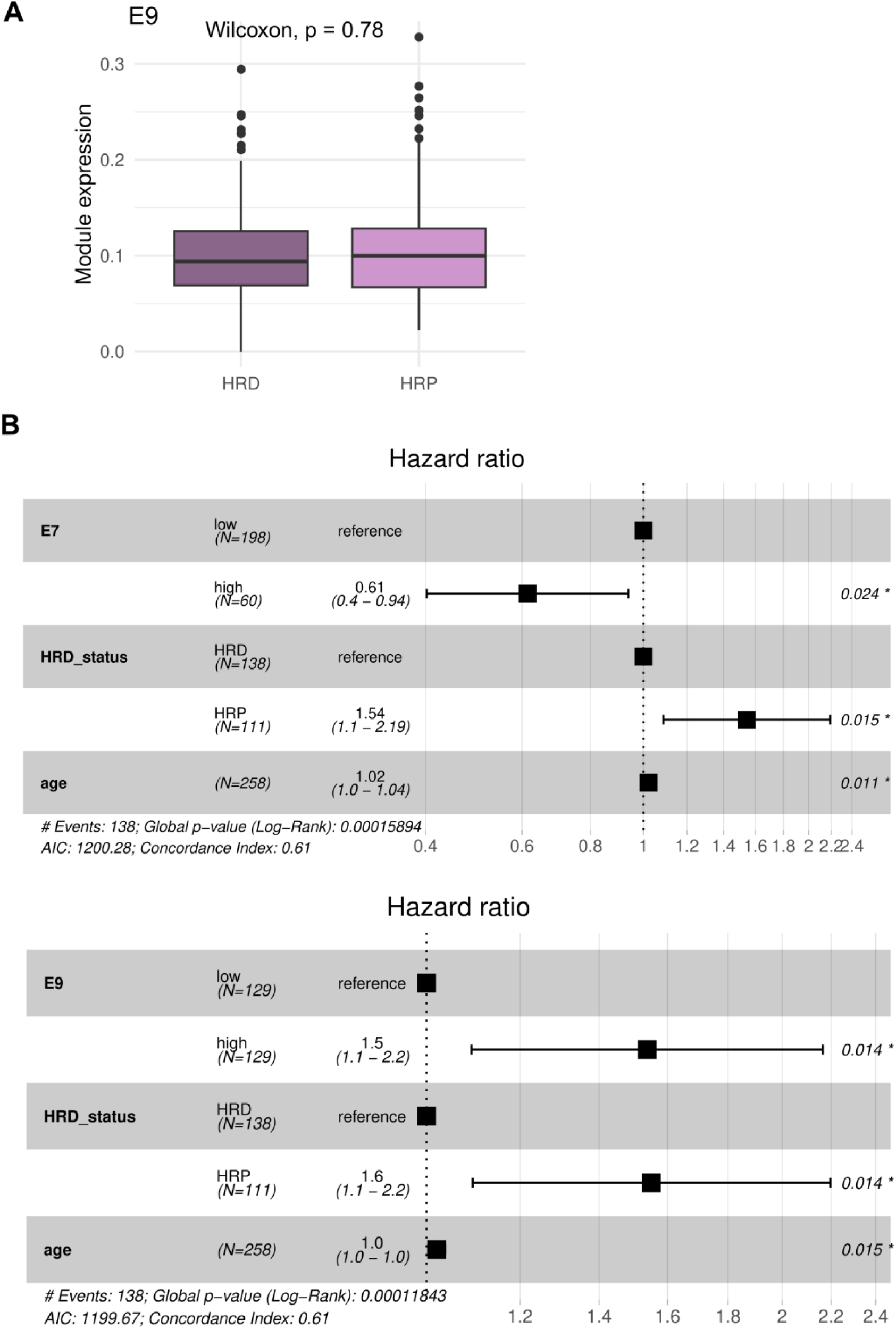
E9 state in HRD and HRP patients, and hazard ratios from multivariate cox regression analysis. (**A**) Box plot showing E9 module activity in homologous recombination deficient (HRD) (n = 138) and homologous recombination proficient (HRP) patients (n = 111) (Wilcoxon rank-sum test, P = 0.78). (**B**) Forest plot showing the results of the multivariate Cox proportional hazard model (hazard ratios, their confidence intervals and *P* values). Modules E7 (above) and E9 (below) were tested whether overall survival (OS) (n = 231 patients) relates them to HRD status or age at diagnosis in the TCGA cohort.

**Fig. S7.**
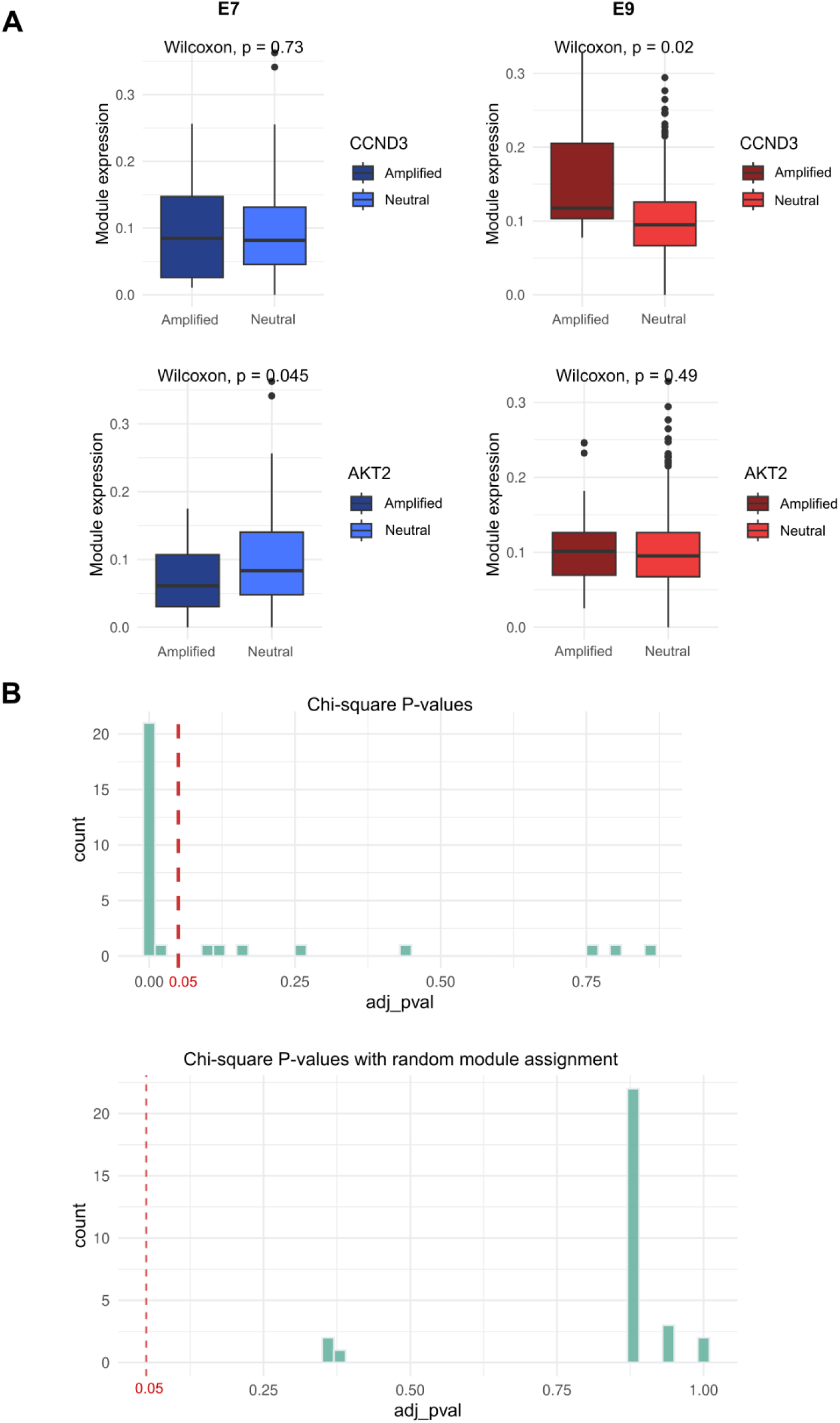
Genetic associations of the modules. (**A**) Box plots showing E7 and E9 module activities in *CCND3* amplified (n = 10) and neutral (n = 238) patients in the TCGA cohort (above). Box plots showing E7 and E9 module activities in *AKT2* amplified (n = 25) and neutral (n = 223) patients in the TCGA cohort (below). (**B**) Histogram of FDR-adjusted *P*-values from Chi-square test between the E7 and E9 gene modules, and subclones in 30 patients (above). Histogram of FDR-adjusted *P*-values from Chi-square test between groups of the same size as the E7 and E9 groups with randomly assigned cells, and subclones in 30 patients (below). Comparison was done for subclone groups having more than 10 cells.

**Fig. S8.**
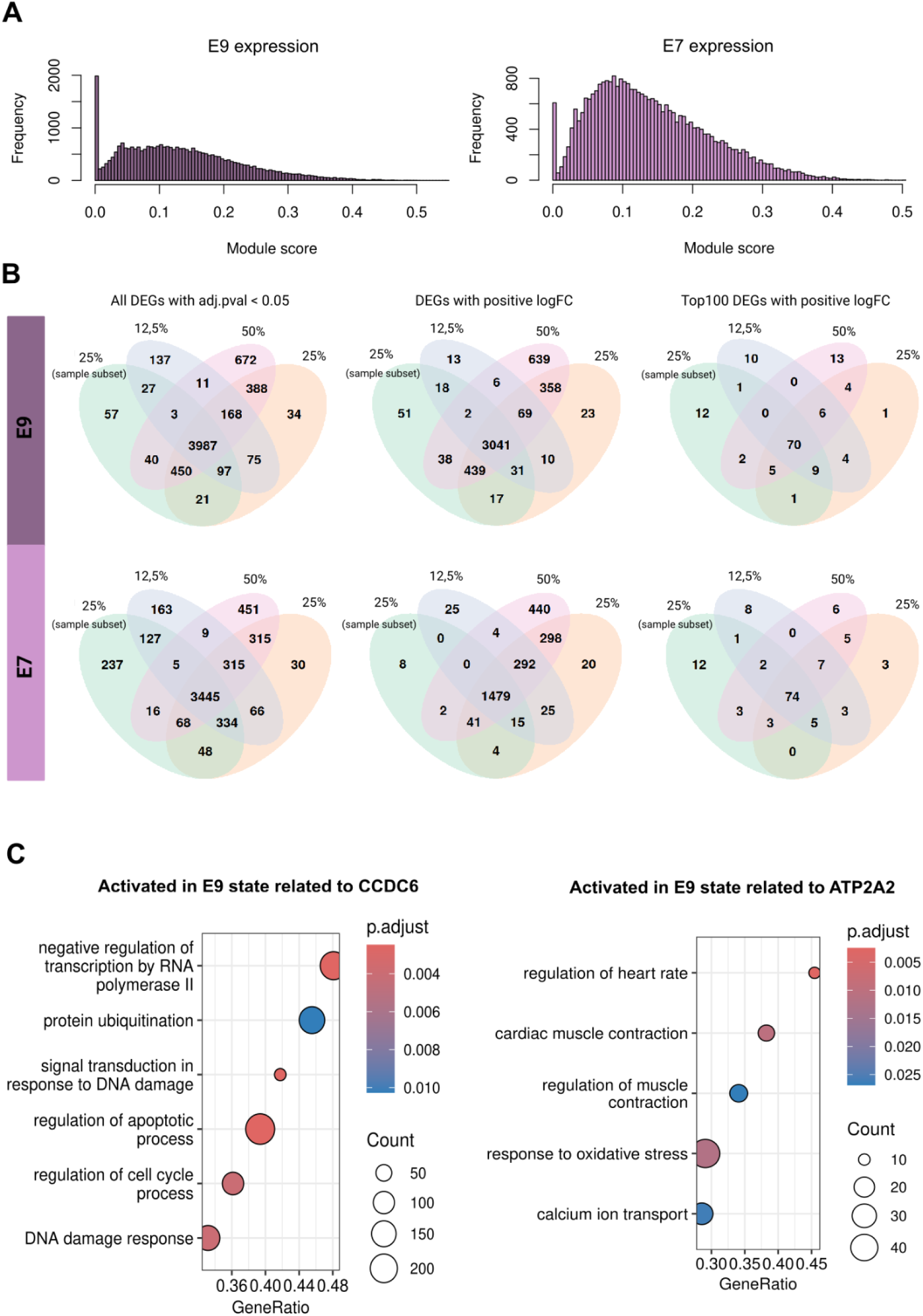
Module expression distributions, justification for threshold 25% for module high cells and biology of E9 driver genes. (**A**) Histograms of distributions of E7 (left) and E9 (right) module activities in cancer cells calculated with UCell. (**B**) Venn diagrams illustrating overlapping differentially expressed genes (DEGs) in E9 (above) and E7 (below) module signatures when 12.5%, 25%, and 50% of cells with the highest module score were used to define cells in the module. We used a 25% threshold to define cells in the module and further subsetted them by averaging cell counts across all samples to reduce sample-specific signals. Overlapping DEGs are shown for all DEGs with FDR-adjusted *P* values < 0.05, for DEGs with positive logFC and for top100 DEGs with positive logFC. (**C**) GSEA scores for GO terms (with Bonferroni-adjusted *P* < 0.05) were calculated between E7 and E9 modules using 25% of cancer cells per sample showing the highest activity of the module. The GO terms related to known biological processes of driver genes are shown in the plot.

**Fig. S9.**
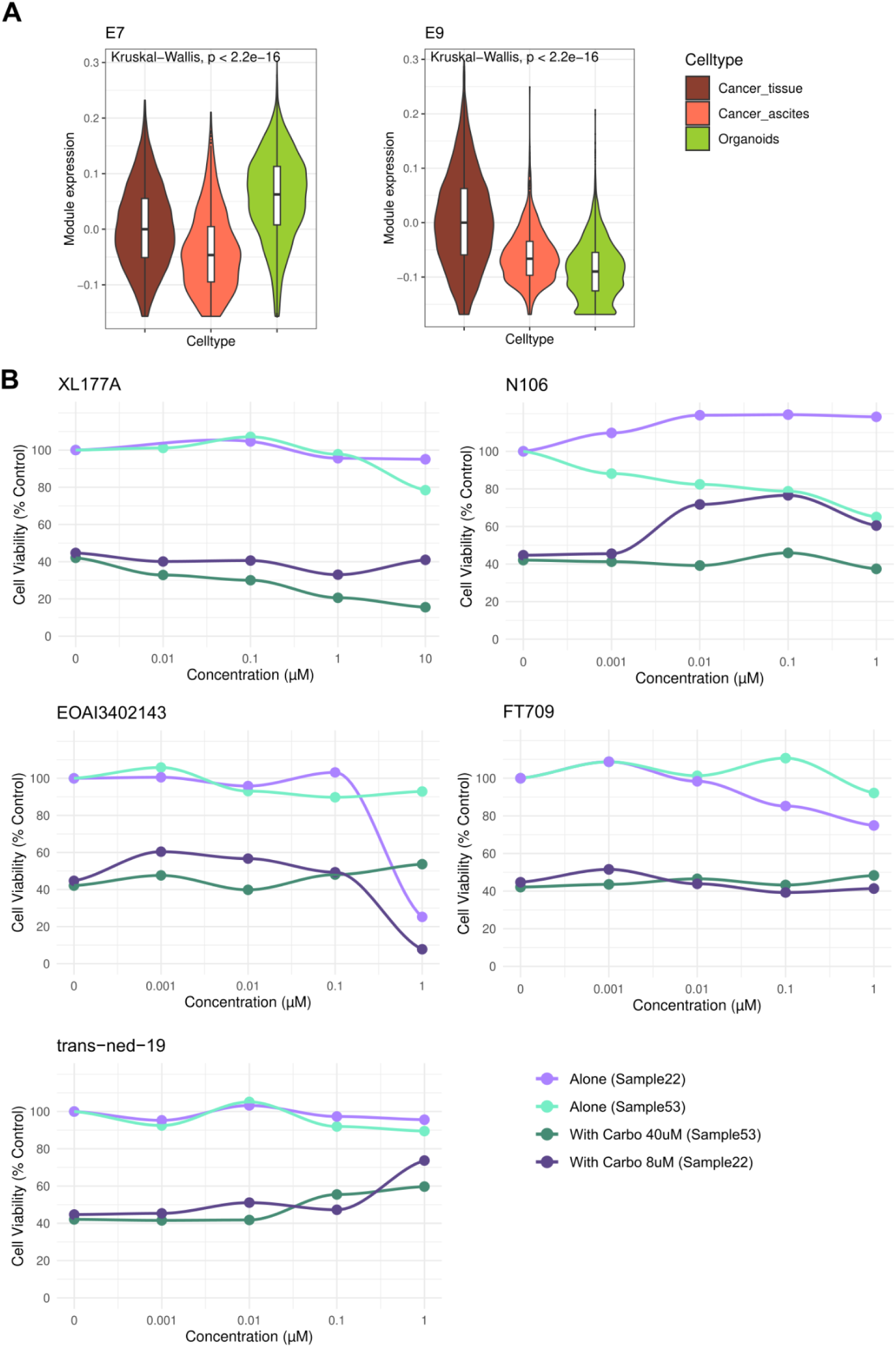
Module activities in organoids compared to tumor tissue, and cell viability results for drug combinations. (**A**) Violin plots showing module activity differences (Kruskal-Wallis test) between tumor tissue (n = 5 767), ascites (n = 1 377) and organoid (n = 5 743) samples at single-cell level. (**B**) Dose-response curves of candidate drugs alone and in combination with carboplatin. Drugs were tested at four different concentrations: XL177A (0.01-10 µM) and the other drugs (0.001-1 µM). Carboplatin was used at 8 µM for sample22 organoid and 40 µM for sample53 organoid. The data shown is normalized to the positive and negative controls.

**Fig. S10.**
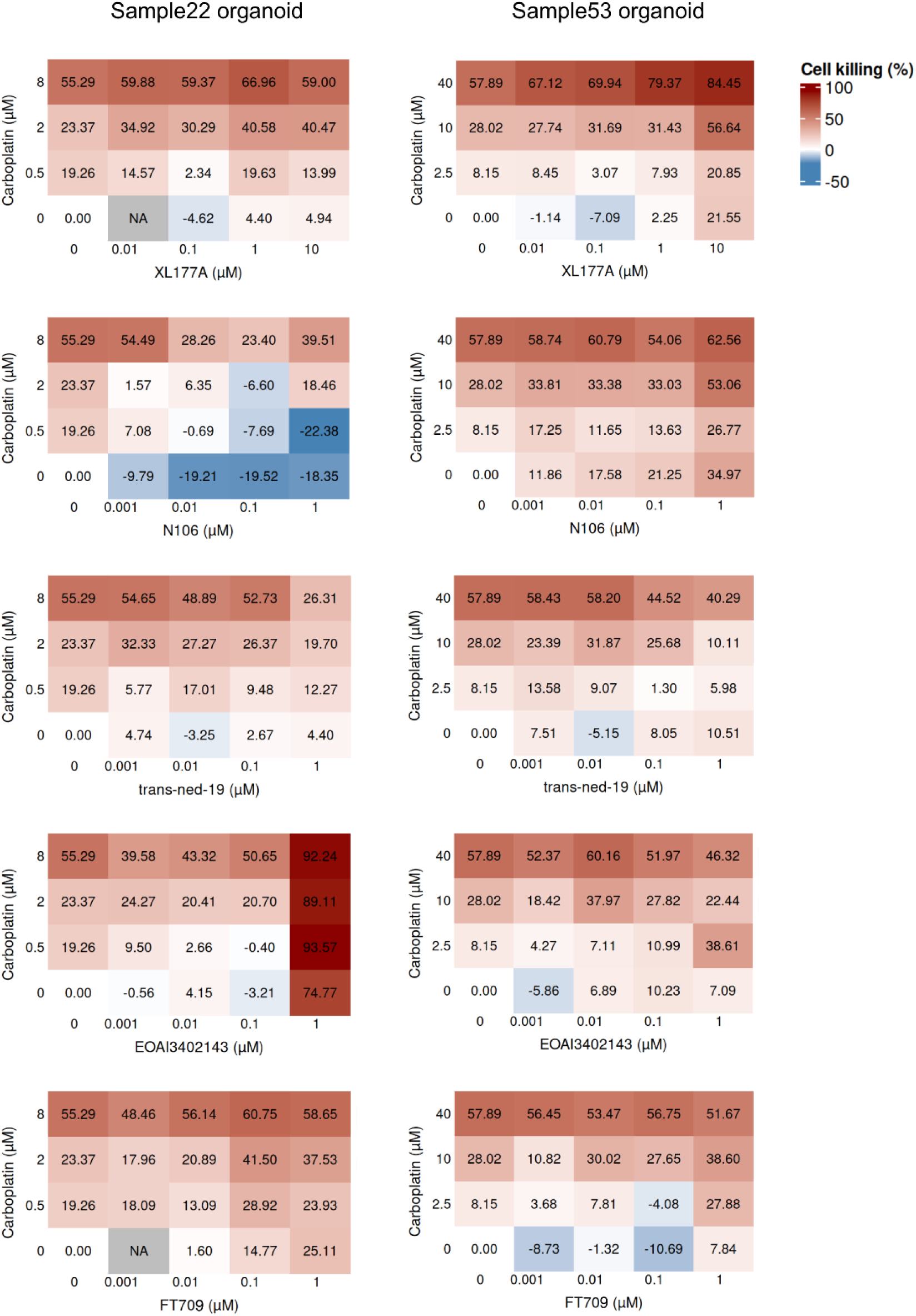
Drug screening results for tested drugs. Dose-response matrices for drug combinations, where organoid cultures were first treated with drugs at concentrations indicated on the horizontal axis, followed by carboplatin treatment at concentrations indicated on the vertical axis.

**Fig. S11.**
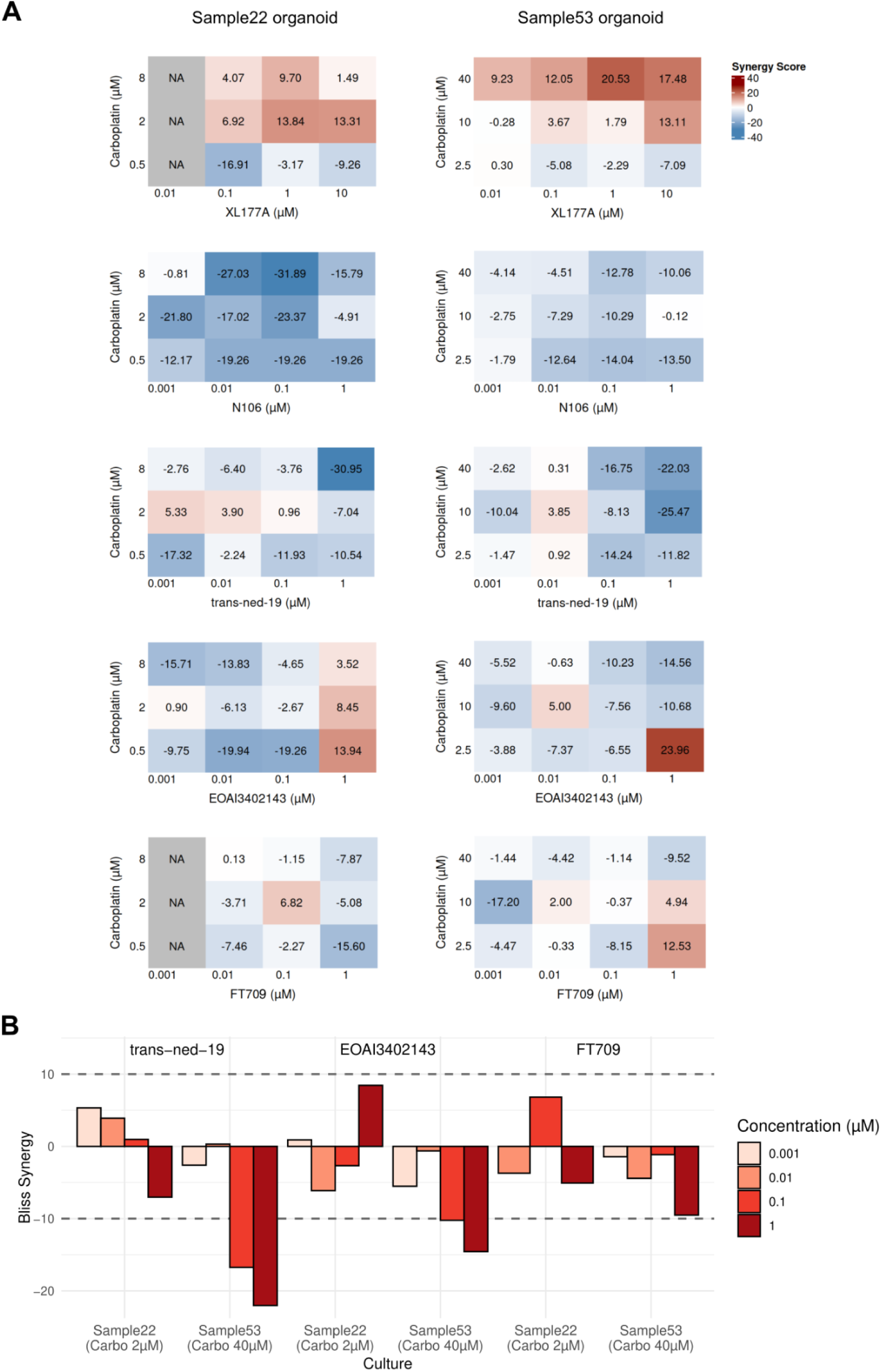
Drug synergies with carboplatin. (**A**) Synergy matrices for the drugs combined with carboplatin. Synergy scores were calculated using Bliss independence model. (**B**) Bliss synergy scores for other drugs at carboplatin concentration 2 µM in sample22 organoid and 40 µM in sample53 organoid. A synergy score greater than 10 was used as the threshold for synergistic interactions, while a score less than −10 indicated antagonistic interactions.

**Fig. S12.**
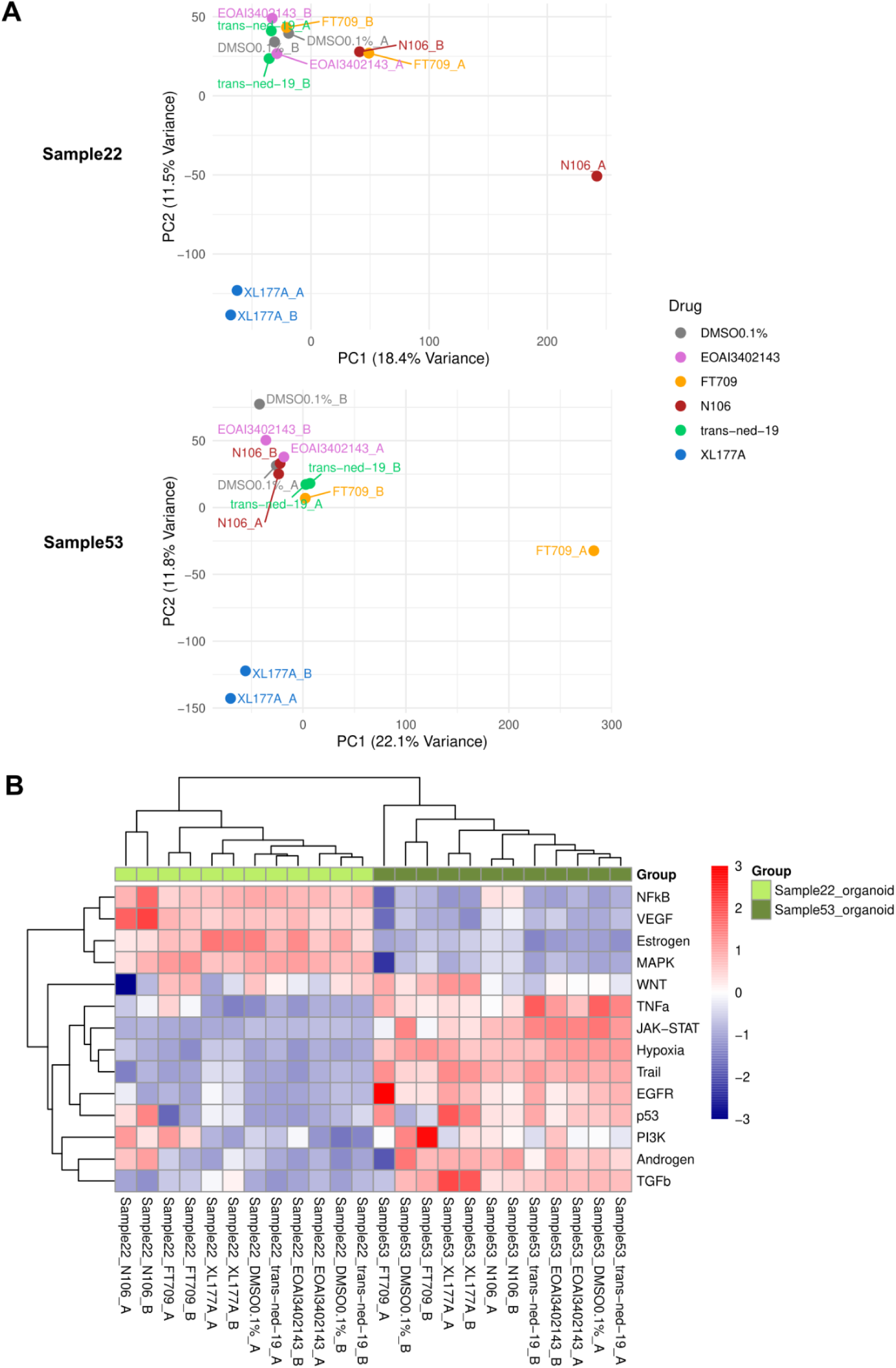
Transcriptomes of treated organoid samples. (**A**) PCA results on the gene expression data of treated and control organoid samples from sample22 and sample53. (**B**) A heatmap showing pathway activities, inferred using PROGENy, in treated and control organoid samples from sample22 and sample53. All the treatments have replicates A and B.

## Supplementary tables

Table S1: Patient cohort clinical data

Table S2: Sample-specific metadata

Table S3: Characteristics of gene modules

Table S4: Genomic comparison of the modules in the TCGA cohort

Table S5: Candidate compounds and associated targets

## Notes

### Competing Interest Statement

The authors have declared no competing interest.

