## Supplementary tables for "Conserved cell state dynamics reveal targetable resistance patterns in ovarian high-grade serous carcinoma"

| PatientID | Sample | Tissue_site | Stage | Age_at_Diagnosis | Published |
|---|---|---|---|---|---|
| Patient1 | sample1 | Tubules | IVA | 57 | Häkkinen_et_al_Bioinformatics |
| Patient2 | sample2 | Ascites | IIIC | 72 | Häkkinen_et_al_Bioinformatics |
| Patient3 | sample3 | Omentum | IVB | 81 | Häkkinen_et_al_Bioinformatics |
| Patient4 | sample4 | Omentum | IVB | 77 | Launonen_et_al_CancerCell |
| Patient5 | sample5 | Peritoneum | IIIC | 60 | Launonen_et_al_CancerCell |
| Patient6 | sample10 | Omentum | IIIC | 78 | Launonen_et_al_CancerCell |
| Patient7 | sample6 | Omentum | IIIC | 74 | Launonen_et_al_CancerCell |
| Patient8 | sample7 | Peritoneum | IVB | 68 | Launonen_et_al_CancerCell |
| Patient9 | sample8 | Omentum | IIIC | 75 | Hippen_et_al_PlosCompBiol |
| Patient10 | sample9 | Ovary | IIIC | 75 | Hippen_et_al_PlosCompBiol |
| Patient11 | sample11 | Ascites | IVA | 67 | Häkkinen_et_al_Bioinformatics |
| Patient12 | sample12 | Mesenterium | IIIC | 77 | Launonen_et_al_CancerCell |
| Patient13 | sample13 | Peritoneum | IIIC | 74 | Häkkinen_et_al_Bioinformatics |
| Patient14 | sample14 | Mesenterium | IIIC | 70 | no |
| Patient15 | sample15 | Ovary | IIIC | 78 | Launonen_et_al_CancerCell |
| Patient16 | sample16 | Peritoneum | IIIC | 68 | Zhang_et_al_SciAdv |
| Patient17 | sample17 | Ovary | IIIC | 69 | no |
| Patient18 | sample18 | Omentum | IIIC | 81 | Launonen_et_al_CancerCell |
| Patient19 | sample19 | Omentum | IVA | 54 | Zhang_et_al_SciAdv |
| Patient20 | sample20 | Tumor | IIIC | 63 | Hippen_et_al_PlosCompBiol |
| Patient21 | sample21 | Tumor | IIIC | 62 | Hippen_et_al_PlosCompBiol |
| Patient22 | sample22 | Omentum | IIIC | 62 | Zhang_et_al_SciAdv |
| Patient23 | sample23;<br>sample24 | Adnexa;<br>Ascites | IIIC | 74 | Senkowski_et_al_DevCell;<br>Senkowski_et_al_DevCell |
| Patient24 | sample25 | Peritoneum | IVA | 67 | Zhang_et_al_SciAdv |
| Patient25 | sample26 | Peritoneum | IIIC | 62 | Zhang_et_al_SciAdv |
| Patient26 | sample27 | Mesenterium | IVA | 64 | Zhang_et_al_SciAdv |
| Patient27 | sample28 | Peritoneum | IIIC | 68 | no |
| Patient28 | sample29 | Peritoneum | IVB | 72 | Launonen_et_al_CancerCell |
| Patient29 | sample53 | Peritoneum | IIIC | 60 | no |
| Patient30 | sample30 | Peritoneum | IVA | 72 | Zhang_et_al_SciAdv |
| Patient31 | sample31 | Peritoneum | IVA | 73 | Zhang_et_al_SciAdv |
| Patient32 | sample32 | Omentum | IVA | 78 | Zhang_et_al_SciAdv |
| Patient33 | sample33 | Omentum | IIIC | 79 | Senkowski_et_al_DevCell |
| Patient34 | sample34 | Peritoneum | IIIC | 72 | Launonen_et_al_CancerCell |
| Patient35 | sample35 | Peritoneum | IVB | 67 | Zhang_et_al_SciAdv |

| | | | | | |
|---|---|---|---|---|---|
| Patient36 | sample36 | Omentum | IIIC | 71 | Launonen_et_al_CancerCell |
| Patient37 | sample37 | Omentum | IIIC | 75 | Launonen_et_al_CancerCell |
| Patient38 | sample38 | Peritoneum | IIIC | 85 | no |
| Patient39 | sample39 | Omentum | IVA | 74 | Zhang_et_al_SciAdv |
| Patient40 | sample40; sample41 | Omentum | IIIC | 68 | no |
| Patient41 | sample42 | Omentum | IIIC | 77 | Launonen_et_al_CancerCell |
| Patient42 | sample43 | Ovary | IIIC | 73 | no |
| Patient43 | sample44 | Omentum | IVB | 55 | no |
| Patient44 | sample45 | Peritoneum | IIIC | 78 | Launonen_et_al_CancerCell |
| Patient45 | sample46 | Omentum | IIIC | 73 | no |
| Patient46 | sample47 | Ascites | IIIC | 73 | no |
| Patient47 | sample48 | Ascites | IIIB | 67 | no |
| Patient48 | sample49 | Ascites | IIIC | 68 | Launonen_et_al_CancerCell |
| Patient49 | sample50 | Omentum | IIIC | 70 | no |
| Patient50 | sample51 | Omentum | IIIC | 75 | no |
| Patient51 | sample52 | Omentum | IVA | 68 | no |
| Patient52 | control1 | Fimbria | | | no |
| Patient53 | control2 | Fimbria | | | Perkiö_et_al_SciRep |
| Patient54 | control3 | Tube | | | Perkiö_et_al_SciRep |
| Patient55 | control4 | Fimbria | | | Perkiö_et_al_SciRep |
| Patient56 | control5 | Fimbria | | | Perkiö_et_al_SciRep |
| Patient57 | control6 | Tube | | | Perkiö_et_al_SciRep |
| Patient58 | control7 | Fimbria | | | no |

**Table S1. Patient cohort clinical data.**

| SampleID | Cancer | NormalFTE _secretory | NormalFTE _cilial | Immune | Stroma | Organoid _RNAseq |
|---|---|---|---|---|---|---|
| sample1 | 211 | | | 189 | 240 | no |
| sample2 | 9 | | | 691 | 0 | no |
| sample3 | 198 | | | 127 | 110 | no |
| sample4 | 50 | | | 433 | 51 | no |
| sample5 | 52 | | | 195 | 9 | no |
| sample6 | 224 | | | 721 | 32 | no |
| sample7 | 51 | | | 185 | 42 | no |
| sample8 | 7 | | | 85 | 348 | no |
| sample9 | 245 | | | 79 | 149 | no |
| sample10 | 65 | | | 282 | 128 | no |
| sample11 | 673 | | | 2 | 0 | no |
| sample12 | 2631 | | | 63 | 2 | no |
| sample13 | 157 | | | 1231 | 1003 | no |
| sample14 | 2686 | | | 175 | 48 | no |
| sample15 | 877 | | | 1422 | 127 | no |
| sample16 | 777 | | | 479 | 224 | no |
| sample17 | 831 | | | 943 | 865 | no |
| sample18 | 3533 | | | 610 | 324 | no |
| sample19 | 1701 | | | 710 | 521 | no |
| sample20 | 46 | | | 835 | 413 | no |
| sample21 | 131 | | | 562 | 2079 | no |
| sample22 | 186 | | | 1717 | 322 | yes |
| sample23 | 775 | | | 113 | 2288 | no |
| sample24 | 511 | | | 3085 | 4 | no |
| sample25 | 178 | | | 3238 | 406 | no |
| sample26 | 200 | | | 1194 | 361 | no |
| sample27 | 200 | | | 1650 | 1207 | no |
| sample28 | 499 | | | 1023 | 233 | no |
| sample29 | 816 | | | 1601 | 94 | no |
| sample30 | 3545 | | | 856 | 67 | no |
| sample31 | 591 | | | 3152 | 416 | no |
| sample32 | 400 | | | 856 | 583 | no |
| sample33 | 509 | | | 4949 | 447 | no |
| sample34 | 66 | | | 1800 | 331 | no |
| sample35 | 49 | | | 1712 | 640 | no |

| | | | | | | |
|---|---|---|---|---|---|---|
| sample36 | 7 | | | 5144 | 206 | no |
| sample37 | 548 | | | 669 | 265 | no |
| sample38 | 425 | | | 1727 | 70 | no |
| sample39 | 49 | | | 2277 | 41 | no |
| sample40 | 540 | | | 536 | 41 | no |
| sample41 | 228 | | | 2319 | 162 | no |
| sample42 | 223 | | | 2286 | 140 | no |
| sample43 | 947 | | | 2437 | 26 | no |
| sample44 | 156 | | | 74 | 41 | no |
| sample45 | 78 | | | 2278 | 8 | no |
| sample46 | 116 | | | 3246 | 51 | no |
| sample47 | 1303 | | | 281 | 1 | no |
| sample48 | 3 | | | 2374 | 162 | no |
| sample49 | 22 | | | 2885 | 6 | no |
| sample50 | 408 | | | 1195 | 32 | no |
| sample51 | 186 | | | 393 | 30 | no |
| sample52 | 1317 | | | 792 | 79 | no |
| sample53 | | | | | | yes |
| control1 | | 252 | 940 | 2270 | 1128 | no |
| control2 | | 143 | 136 | 1879 | 186 | no |
| control3 | | 75 | 139 | 1790 | 634 | no |
| control4 | | 575 | 289 | 2433 | 54 | no |
| control5 | | 41 | 349 | 889 | 1 | no |
| control6 | | 158 | 446 | 2624 | 7 | no |
| control7 | | 7 | 14 | 1001 | 1489 | no |

**Table S2. Sample-specific metadata.**

| Module | Genes |
|---|---|
| N1 | ACSL5, ACSM3, ACSS3, ADGRG2, AFAP1L1, AKAP7, ALDH1A1, ALDH1A2, ALDH6A1, APOBEC3C, APOBEC3F, APOL4, CCDC122, CD22, CDH16, CETN3, CHDH, CLCN3, CLCN5, CLYBL, CRTAC1, CRYL1, CYP4V2, CYYR1, ERP27, FAM13A, FAXDC2, FERMT1, FGGY, FOLH1, FRY, GJB1, HNF1B, IKZF4, IMPG2, KCNMB2, KCNQ1, KIF12, LCN12, LRG1, LTC4S, MAG, MAGEH1, MAOA, MCF2L, MLPH, MMP28, MUC15, NDP, NOSTRIN, NPNT, NRGN, NXF3, OBP2A, PAK3, PAX2, PGA5, PGM5, PLA2R1, PLEKHH2, PLLP, PLSCR4, POF1B, PRSS33, PTCH1, PTPN3, PYROXD2, SCGB1A1, SCGB3A1, SFXN2, SIAE, SLC15A2, SLC16A9, SLC40A1, SLC7A8, SMIM10, STEAP2, SYK, TBX2, THNSL2, THRA, TLR5, TMEM45B, TUNAR, UGT8, VSIG2, WNK2, ZG16B |
| N2 | ABO, AGR2, AGR3, ANAPC4, BBS4, BTG2, C20orf85, C9orf24, CCDC160, CCDC170, CCDC191, CDKL1, CFAP43, CFAP44, CFAP53, CFAP69, CFAP70, CLUAP1, CYP2U1, DMD, DNAJB13, DRC3, DYNC2H1, DYNLRB2, DZIP3, EPHX2, FAM13C, FAM47E, FILIP1, FOXJ1, GEMIN8, GJA4, GRAMD1C, GSTA1, HYDIN, IDNK, IFT74, IFT88, IL20RA, INTU, KIF27, LIAS, LYRM9, MAOB, MIA2, NEIL1, NME5, NPDC1, NUDT6, NUDT7, PACRG, PALMD, PIFO, PIGV, PLCH1, POLI, PRUNE2, PTPRN2, RASSF6, RFX3, SLC16A11, SMIM6, SORBS2, SPATA18, SPATA7, SPEF2, SPIRE2, SSBP2, STK33, TFF3, TSPAN2, UBXN10, USP51, ZC2HC1A, ZC3H6 |
| N3 | ADAM28, ANKRD18A, AR, ARHGAP26, ARHGEF10, BMPR1B, BNC2, BTBD8, BTNL9, CCDC171, CCSER1, CD200, CDH12, CHM, CNGA1, CNTN4, COL4A5, DIAPH2, DPH6, ESR1, FAM111A, GAMT, GKAP1, GNB5, GPHN, GREB1, GSAP, GUCY1A2, HRH1, INSR, INTS6L, ITPR2, KCNJ2, L3MBTL4, LGR5, LONRF2, MAGI1, MAML3, MBNL3, MBOAT1, MEGF11, MEGF9, MLLT3, MPPED2, MST1, NHS, NRXN3, PDE3A, PIK3C2G, PLA2G4A, PPP2R2B, PRDM5, PREX2, PTPRM, RASSF9, RERG, RGS7BP, RSPO1, SCAPER, SEMA4A, SERPINA5, SETBP1, SLC7A2, SLITRK5, SORL1, SOX6, TDRD3, TEC, THSD4, TNFRSF19, VWA8, ZBTB20, ZFPM2, ZMAT1, ZNF334 |
| N4 | ADCYAP1R1, C2orf88, CLDN22, CRISP2, CRISP3, DEPTOR, LENG9, NMNAT3, NXNL2, OVGP1, PGR, PKHD1L1, RIMS2, SLC27A6, SOX3, SPDEF, TMEM220, TSPAN8, TTYH1 |
| C1 | AAGAB, ABRACL, ACOT7, ALG3, ANLN, ARHGAP11A, ASF1B, ASPM, AURKA, BIRC5, CCNB2, CCNE1, CCSAP, CDC20, CDCA5, CDCA8, CDK1, CDKN2D, CDKN3, CENPA, CENPF, CENPU, CENPW, CEP55, CHEK1, CLSPN, CTPS1, DBN1, DDX39A, DIAPH3, DNMT1, E2F1, ECE2, ECT2, EIF4EBP1, FAM83D, FANCI, FOXM1, GPN1, HRK, ICMT, KIF23, KIFC1, KNSTRN, KPNA2, KRT16, MAGOHB, MCM4, MTCH2, MYBL2, NDC1, NDUFAF6, PDCD2L, PGP, PI3, PLAU, PLK1, PRAME, PRC1, PTTG1, RACGAP1, RAD51AP1, SGO1, SLC5A6, STMN1, TK1, TPX2, TSEN15, UBE2C, UCHL1, USP39, USP5, XDH, ZNF354A, ZWINT |
| C2 | ATG16L1, B3GALNT2, BAIAP2L1, BRAP, CCNYL1, CDKN2A, CHKA, COMMD2, COMMD7, DEDD, DPH2, DPH3, DUSP12, DYRK2, E2F3, EFHD1, ENO2, FIGN, GAL, GCLM, GMEB2, GMFB, GOLGA7, GORASP2, HES6, IMPDH1, KIF1A, LAGE3, MBOAT2, MFSD5, MFSD9, MTHFD1L, MYO10, NGEF, NRAS, NXPH4, OSGIN2, POP7, PROSER2, RAB22A, RABIF, RAD1, |

| | |
|---|---|
| | RNF2, S100A2, SFN, SMIM10L1, TAF13, TEAD4, TFAP2A, TIMM17A, UBE2F, WRNIP1, YTHDF1, ZNF35, ZNF518B, ZNF639 |
| C3 | ANKRD52, ARL8A, ATG13, ATG9A, AXIN1, B4GALT2, C1orf198, C8orf76, CDR2L, CLPTM1L, DDA1, DENND4B, DNAJC5, DVL3, ECE1, EPHX4, FASN, FBXO45, GIPC1, HGS, HSF1, HYOU1, L1CAM, LRFN4, LRRC8D, MCU, NPLOC4, PKN1, PLXNA1, RNF220, SLC52A2, SLC6A8, SMG5, SQLE, STRN4, TFPT, TMEM132A, TMPRSS4, TNFRSF21, TRIM11, WSB2, YEATS2, YKT6, ZC3H3, ZDHHC9, ZMYND19, ZNF598, ZNF777 |
| C4 | ABCF3, ADRM1, CEBPG, CNOT11, EIF1AD, HSPA14, MRGBP, NANP, NIPA2, PCBP4, PROSER1, RNASEH1, RNF114, RRP36, SCNM1, SEC61A2, SLC35B1, SPECC1, TIMM23, TMEM44, TP53RK, TPD52L2, TRMT6, TSN, TUFT1, WDR53, ZNF48 |
| C5 | ACER3, ADIPOR1, BAK1, MCUR1, PGAM5, PRADC1, RPP25, RUSC1, SLC25A39, TMEM183A, TSPAN17, TUSC2 |
| C6 | ASNS, DDX49, FLAD1, MED8, MTHFD2, PRELID3B, SCAMP3, SIKE1, STC2, TMEM38A, TRIB3 |
| E1 | AAMDC, ABHD11, ADGRG1, AGRN, AHNAK2, AIG1, ALKBH2, AMN, AMOTL2, ANXA3, ASAP2, ASS1, ATN1, ATP6V1B1, BACE2, BAIAP2, BCAM, BCL2L1, BCL2L12, BCL9, BHLHE41, BIK, C19orf33, C6orf132, C8orf82, CAPN1, CAPS, CC2D1A, CCNB1IP1, CD47, CD9, CDC42EP1, CDC42EP4, CDS1, CELSR2, CHMP4C, CITED4, CLDN3, CLDN4, CLDN7, CLMN, CLU, COMTD1, CP, CRABP2, CRACR2B, CRB3, CRIP2, CTXN1, CXCL17, DAG1, DAPL1, DDAH2, DDR1, DEFB1, DHCR24, DHCR7, DOK5, EBPL, EDN1, EFNA1, EMID1, EMX2, EPPK1, EPS8L1, EPS8L2, ERBB2, ERBB3, ESRP1, FAIM, FAM107A, FAM136A, FAM3B, FAM83H, FBXO2, FGF18, FLNB, FOXP4, FRAT2, FXYD3, GIPC1, GMDS, GMPR, GPRC5C, GRB7, HDAC1, HDAC7, HOOK2, HSBP1L1, IFT172, ILVBL, IQCK, ISYNA1, ITGA3, ITGB4, ITPR3, JUP, KAZN, KIAA0040, KIAA1522, KIF9, KLK10, KLK11, KLK6, KLK7, KLK8, KRT17, KRT18, KRT19, KRT23, KRT7, KRT8, LAD1, LAMA3, LAMA5, LAMB3, LCN2, LEMD1, LLGL2, LMO7, LTBP3, LYPD1, LYPD6B, MACC1, MAL2, MARK2, METRN, MFSD3, MISP, MPZL2, MTA1, MUC1, MUC16, MUC4, MYH14, MYO5B, NECTIN4, NELFA, NFKBIL1, NOL4L, NPR1, NR2C2AP, NR2F6, NSG1, NT5C3B, NUDT14, OBSL1, PBX2, PCDH7, PDE4C, PDLIM1, PDZK1IP1, PERP, PIP4K2C, PKP3, PLCD3, PLXNB1, PLXNB2, PNOC, PPL, PPP1R13L, PPP1R16A, PPP1R1B, PRRG4, PRSS22, PRSS8, PTGS1, PTP4A3, PTPRF, QTRT1, RAB25, RAB5B, RASSF7, RBP1, RCC1, REC8, RFK, RGL2, RHOV, RIC3, RIPK4, RNF39, RTKN, S100A1, S100A13, S100A14, SCARA3, SCGB1D2, SCGB2A1, SCRIB, SCRN1, SEMA3B, SH2D3A, SH3GLB2, SLC25A23, SLC25A29, SLC34A2, SLC39A4, SLC40A1, SLC44A2, SLC44A4, SLC48A1, SLPI, SMARCA4, SMARCD3, SORT1, SOX17, SOX9, SPINT1, ST14, STON2, STRBP, STXBP6, SUDS3, SYNGR2, TACC2, TACSTD2, TBC1D17, TCEA3, THEM6, TMEM134, TMEM205, TMEM238, TMEM30B, TMOD1, TMPRSS3, TNNT1, TP53, TRIM16, TRIM29, TRIP6, TSPAN12, TTC9, TYW3, WFDC2, WRNIP1, WT1, YIPF2, ZFAND1, ZNF165, ZNF3, ZNF592, ZNF664, ZNF768 |
| E2 | ADNP, AHCY, ALDH3A2, ARHGAP29, ARHGEF12, ASRGL1, BAIAP2L1, BICD1, BTBD3, C2orf88, CCDC112, CCDC6, CCDC85C, CCNA1, CCNC, CD24, CD2AP, CDH1, CDH6, CEP70, CHD3, CLDN1, CLDN10, CLGN, COBL, CRIM1, |

| | |
|---|---|
| | CSRP1, CTNND1, CXADR, DCDC2, DNAH14, ECSIT, ECT2, EFNB2, EHF, ELF3, EPB41L5, EPN2, ESR1, EYA2, F11R, F2RL1, FARP1, FGD4, FGFR2, FNBP1L, GALNT3, GCLC, GTF2I, HACD2, HOMER2, HOOK1, ID4, IGF2BP2, INTS3, ITGA6, ITGB6, KCNK6, KLF5, KMT5B, LAMC2, LARP1, LDLR, LRP6, MACROD2, MAN1A2, MAP3K13, MAP7, MEIS1, MMP7, MTMR12, MYEF2, MYH10, MYO6, MYOF, NAB1, NET1, NFIB, NFKBID, NUAK2, NUDT15, NUP88, OVOL2, PARD3, PATJ, PAWR, PAX8, PBX1, PEG10, PELP1, PKN2, PKP2, PKP4, PLA2G12A, PLAGL2, PLEKHA5, PODXL, PRKCI, PROSER2, PTPN14, PTPRK, PTRHD1, RAB11FIP1, RALBP1, RALGPS2, RBBP8, RBM47, RHPN2, RIMKLB, ROCK2, RPRD1A, RTN3, SCNN1A, SENP2, SGPP2, SLC12A2, SLC25A36, SLC7A1, SPIN1, SRGAP3, SUN1, SYNJ2BP, TBL1XR1, TC2N, TEAD1, TFAP2C, TGIF2, TPD52, TSPAN13, UACA, UGT2B7, USP6NL, VTCN1, WDR73, WWC1, XPR1, YAP1, YOD1, ZC3H14, ZMYND8, ZNF217, ZNF322, ZNF33A |
| E3 | ACYP1, ALG8, ARL2, ARMC10, ATIC, BEX3, CCDC34, CDKN2AIPNL, CHCHD6, COA3, COA4, COA5, DMKN, DNAJC19, DPCD, DPY30, ECHS1, EPCAM, FDPS, FUNDC1, GCSH, GGCT, HDDC2, HMGN3, HSPBP1, IFT22, IFT57, LAPTM4B, LSM4, MAGEF1, METTL5, MGMT, MMAB, MORN2, MRPL24, MRPL3, MRPL48, MRPS25, MRPS26, MRPS33, MTX2, NAXE, NDUFV1, NIT2, NTPCR, NUDCD2, PEMT, PFDN6, PFN2, PHGDH, POP5, PPA2, PRDX3, PRMT5, PSAT1, PXMP2, QPRT, RCN2, RFXANK, RIDA, RUVBL1, RUVBL2, SCCPDH, SCOC, SDHAF3, SH3YL1, SLC25A11, SPAG16, SSRP1, STK25, SUCLG1, SYNE4, TARBP2, TIMMDC1, TMEM106C, TMEM14A, TMEM223, TMEM41A, TMEM9, TMEM97, TPD52L1, TRAPPC6A, TSEN34, TSPAN6, TUSC3, VPS72, ZCRB1 |
| E4 | AIFM1, ALDH7A1, ATXN7L3B, CNOT7, CTBP2, EBAG9, EHMT2, GLO1, HDAC2, KCTD1, MECOM, MED29, MNAT1, MRPS9, MRS2, MTA3, MTIF2, NDUFA5, ORC4, PDHA1, PRPF6, SLC39A10, TOM1L1, ZNF518A, ZSCAN18 |
| E5 | ANAPC15, AP1M2, ECI1, HINT2, ITPA, LDOC1, MANBAL, NAA38, NTHL1, P4HTM, PNKD, PTGES2, PTOV1, PTRH1, SSBP4, TCEA2, TM7SF2, TMEM125, TMEM141, TRPT1, TSTD1, UQCC3 |
| E6 | ATP6V0E2, ATPAF1, BEX2, BMP7, CYB561, ESPN, FKBP4, IGSF8, IMMP1L, MALSU1, NT5DC2, NUPR2, OGFOD3, PAK4, PPP5C, REPIN1, SELENBP1, SLC25A4, TM7SF3, TSPAN1, UCKL1 |
| E7 | CKS1B, DCTPP1, DNPH1, ERI3, EXOSC5, FIBP, GSTO2, KRTCAP3, MACROD1, MRPL11, MYL6B, NDUFB5, NHP2, NIPSNAP1, OLA1, PAFAH1B3, PAICS, PCBD1, POLD2, TATDN1, TRAP1 |
| E8 | AGPAT2, APOA1, ARRDC1, CNFN, FLOT1, FOLR1, GALE, KCNK15, LSR, MGST2, MSLN, PLPP2, PON2, PPP1R35, SMIM22, SNCG, SPINT2, TMC4, TSPAN15, VWA1 |
| E9 | ANKRD33B, DSP, EPHA2, IRF6, ITPKC, OCLN, PARD6B, PIP5K1A, PPP2R3A, RAB3IP, SGMS2, TES, TJP1, TUFT1, UBE2H |

**Table S3. Characteristics of gene modules.**

| Gene | CNstatus | Count | E7_Pval | E7_adjPval | E9_Pval | E9_adjPval |
|---|---|---|---|---|---|---|
| AKT2 | AMP | 25 | 0.04542 | 0.3039 | 0.4942 | 0.799328571 |
| AKT3 | AMP | 10 | 0.8307 | 0.9345375 | 0.8908 | 0.8914 |
| CCND2 | AMP | 22 | 0.9442 | 0.9442 | 0.6217 | 0.799328571 |
| CCND3 | AMP | 10 | 0.7307 | 0.9345375 | 0.02036 | 0.18324 |
| CCNE1 | AMP | 49 | 0.1013 | 0.3039 | 0.3357 | 0.799328571 |
| KRAS | AMP | 36 | 0.5616 | 0.9345375 | 0.5348 | 0.799328571 |
| MECOM | AMP | 56 | 0.7807 | 0.9345375 | 0.8914 | 0.8914 |
| MYC | AMP | 61 | 0.09863 | 0.3039 | 0.3361 | 0.799328571 |
| PIK3CA | AMP | 39 | 0.6354 | 0.9345375 | 0.3702 | 0.799328571 |

**Table S4. Genomic comparison of the modules in the TCGA cohort.**

| **Drug** | **Gene** | **Module** | **Action** | **MOA** |
|---|---|---|---|---|
| EOAI3402143 | USP9X, USP24, USP5 | E7, E9 | inhibition | deubiquitinase inhibitor |
| FT709 | USP9X | E7, E9 | inhibition | potent and selective USP9X inhibitor |
| N106 | ATP2A2 | E9 | activation | SERCA2a activator through SUMOylation |
| trans-Ned-19 | LSM12 | E7, E9 | inhibition | potent and selective inhibitor of NAADP receptor |
| XL177A | CCDC6 | E7, E9 | inhibition | USP7 inhibition through a p53-dependent mechanism |

**Table S5. Candidate compounds and associated targets.**
